# Fitness landscapes for species interactions: when do population genetics and adaptive dynamics diverge?

**DOI:** 10.64898/2026.03.17.712462

**Authors:** Kasturi Lele, Lawrence H. Uricchio

## Abstract

Multiple frameworks have been developed to investigate the evolution of species interactions on fitness landscapes, each with unique strengths and weaknesses. These include adaptive dynamics, which uses linear stability analyses to predict eco-evolutionary outcomes resulting from the invasion of rare mutants into a resident population, and population genetics, which mechanistically models finite populations and stochastic processes in finite time. Though there are some known correspondences between these frameworks, it is not clear that they will always result in the same eco-evolutionary outcomes. Moreover, while adaptive dynamics is very powerful for predicting outcomes, it is not always straightforward to relate these predictions to the data generated in experimental evolution studies. Here, we use a data-driven model of microbial species interactions to compare and contrast the predictions of population genetics and adaptive dynamics. We derive expected outcomes for one-species and two-species evolutionary trajectories by using the invasion fitness landscape concept from adaptive dynamics, and then use analytical theory and forward-in-time simulations to set these predictions within the context of population genetic models. In the context of our one-species models, we show that the timescale of evolution depends on mutation supply and effect sizes, when populations are initialized both along and off a trade-off function. For two-species competition models, we show that mutation supply, effect sizes, and asymmetries between competing species result in discrepancies between adaptive dynamics and population genetics, especially in cases where adaptive dynamics predicts stable coexistence. Our study provides insight into the role of finite timescales, mutation supplies and population sizes in the evolution of species interactions, and facilitates further research that leverages the invasion fitness landscape concept within the realm of population genetics.

## Introduction

Interactions between species are major determinants of community structure (HilleRisLambers et al., 2012), and evolutionary processes are likely to play a major role in shaping these interactions (Thompson, 1999; Harmon et al., 2019; Leibold et al., 2022). Evolving species interactions can be conceptualized as a dynamic process occurring on a fitness landscape, as posited by the eco-evolutionary framework known as adaptive dynamics (Brännström et al., 2013). While traditional fitness landscapes are often framed in terms of multiple genotype or phenotype axes in a population, the axes of the fitness landscape in adaptive dynamics compare the current “resident” value of a trait to a potential rare mutant phenotype (Waxman and Gavrilets, 2005). Viewing the fitness landscape through this lens facilitates analyses that can predict evolutionary outcomes from the ecological dynamics between individuals with resident and mutant trait values. Possible outcomes of such models include the existence of stable fixed points in trait space (evolutionarily stable strategies) and diversification or branching (Ferriere and Legendre, 2013).

While some population genetic models have a natural correspondence with adaptive dynamics (Van Cleve, 2023), mechanistic evolutionary models and adaptive dynamics could also generate discrepant results in some circumstances. Adaptive dynamics is based on differential equations, linear approximations, and stability analyses, and hence it is not always straightforward to relate its findings directly to the genetically explicit modeling frameworks typically used in population genetics. Population genetic-based analyses have sometimes come to different conclusions than models based on adaptive dynamics, and have even challenged the need to invoke fitness trade-offs to explain observed patterns of trait diversity (Whitlock, 1996). Research that directly connects population genetics to adaptive dynamics could advance our understanding of the evolution of species interactions by combining mechanistic insights into evolution in finite populations with the ability to predict long-term eco-evolutionary outcomes.

One challenge in connecting these frameworks is that several assumptions made in adaptive dynamics could be in conflict with evolutionary and ecological processes occurring in real populations and in mechanistic population genetic models (Barton and Polechová, 2005). Specifically, mutations are assumed to be of small effect and rare, occurring one at a time in a clonal population (Otto and Day, 2007; Waxman and Gavrilets, 2005). Population sizes are assumed to be effectively infinite, and traits are often assumed to be constrained to trade-off curves (Boots and Haraguchi, 1999; Bowers and White, 2002; White and Bowers, 2005). The potential impact of each of these assumptions can be considered in light of empirical and theoretical evidence garnered from years of study in population genetics and population ecology.

In real populations, finite size will affect the supply of mutations, the probability of interference between selected alleles, the number of fixation events per generation, and the time required to transit between values in trait space (Orr, 2010; Otto and Whitlock, 2013). Though the effects of mutations are often small, some mutations underlying ecologically-relevant traits can have very large fitness effects (Perfeito et al., 2007; Heilbron et al., 2014; Taylor et al., 2019), perhaps invalidating some assumptions of stability analyses performed in adaptive dynamics. And while adaptive dynamics makes strong predictions about dynamical processes and eco-evolutionary outcomes (such as coexistence or branching), it makes relatively fewer claims about quantities that are easy to measure, such as sequence variation or the various paths through trait space that might be taken in a longitudinal experiment.

Assumptions about the existence and nature of trade-offs between traits are also central to adaptive dynamics, as they are throughout many sub-fields of evolution and ecology (Stearns, 1998). In population ecology, trade-offs between growth rate and species’ yield (*r* − *K* trade-offs) are often assumed to be fundamental, and many studies have found evidence for such trade-offs (Jasmin and Zeyl, 2012; Maharjan et al., 2007; Bachmann et al., 2013; Meyer et al., 2015; Moerman et al., 2020, but see also Reding-Roman et al., 2017; Wei and Zhang, 2019; Marshall et al., 2022). Although constraints on species growth and yield must exist (or population sizes would be unbounded), in adaptive dynamics these constraints are specifically conceptualized as curves in trait-space, along which populations can evolve. Seen through the lens of a traditional fitness landscape where the axes represent distinct combinations of trait values that an individual can attain through mutations, this would be equivalent to constraining the evolutionary process to a single dimension through the planar surface of possible trait values. In mechanistic models of evolution with finite population sizes and changing environments, it is possible that populations may evolve to performance values that fall below the constraints imposed by trade-off curves – for example, through the accumulation of deleterious mutations. trade-off curves can therefore also be conceptualized as bounds on performance rather than constrained paths through trait space (Shoval et al., 2012), which might be more consistent with mechanistic models of evolutionary processes. This latter conceptualization of trade-off curves as performance bounds might also correspond more closely to the results of experimental studies that have sought to infer trade-offs from replicate evolutions (Novak et al., 2006), and is consistent with the traditional conceptualization of fitness landscapes.

Although critiques of adaptive dynamics are not new (Spencer and Feldman, 2005), few studies have sought to assess the robustness of adaptive dynamics predictions in the context of mechanistic population genetic simulation models. A potential challenge for any such assessment is that many of the important quantities (mutation rates of effect alleles, effect size distributions, trade-offs and constraints on trait values, and to some extent, population sizes) do not have empirical estimates in most systems. Indeed, it is often challenging simply to characterize species interactions in ways that make useful predictions about population dynamics and community structure, even in the absence of any evolutionary dynamics. Any effort to assess the robustness of adaptive dynamics will therefore benefit from utilization of prior data-driven models of population/community dynamics to the degree possible, while integrating over a large range of parameter values for the unknown quantities.

Among well-established community assembly models, the generalized Lotka-Volterra (gLV) model has been particularly influential, in part due to its simplicity and its ability to explain empirical patterns in a wide variety of systems (van den Berg et al., 2022). In this model, the change in abundance per unit time depends only on the growth rate and the interaction coefficients, which capture the effects of intra- and inter-specific competition on the growth rate. While gLV models do perform relatively poorly in some empirical systems, they have done particularly well for communities with intermediate species richness (3-10 species; see Gonze et al., 2018; Venturelli et al., 2018; Dedrick et al., 2023). Recently, we characterized interactions in a microbial community with intermediate species richness, finding that we could accurately predict community structure and abundance patterns for seven of nine species with a gLV model (Lele et al., 2026). Since gLV models have broad applicability and can explain patterns of diversity in some fast-evolving microbial systems, a better understanding of evolutionary dynamics of species interactions within the gLV context could advance our ability to predict eco-evolutionary outcomes in some natural microbial communities.

In this work, we study the evolution of species interactions in the context of a gLV model informed by empirical parameter estimates (Figure 1). We use the invasion fitness landscape of adaptive dynamics to set baseline expectations for eco-evolutionary outcomes, and we test whether these expectations hold in both single-species and two-species stochastic population genetic simulations. In general, our goal is to better relate adaptive dynamics predictions to measurable experimental quantities, such as rates of differentiation at the sequence and trait levels, and co-occurrence patterns over finite timescales. In the context of single-species evolution, we use population genetic models to ask how the rate of evolution at the trait and sequence levels scales with mutation rates, effect sizes, and population sizes, and we consider how the timescale of evolution may affect the inference of trade-off functions in long-term evolution studies. For multi-species evolutionary models, we consider whether population genetic models with finite population sizes and supplies of beneficial mutations are consistent with coexistence/stability predictions made by adaptive dynamics.

**Figure 1:**
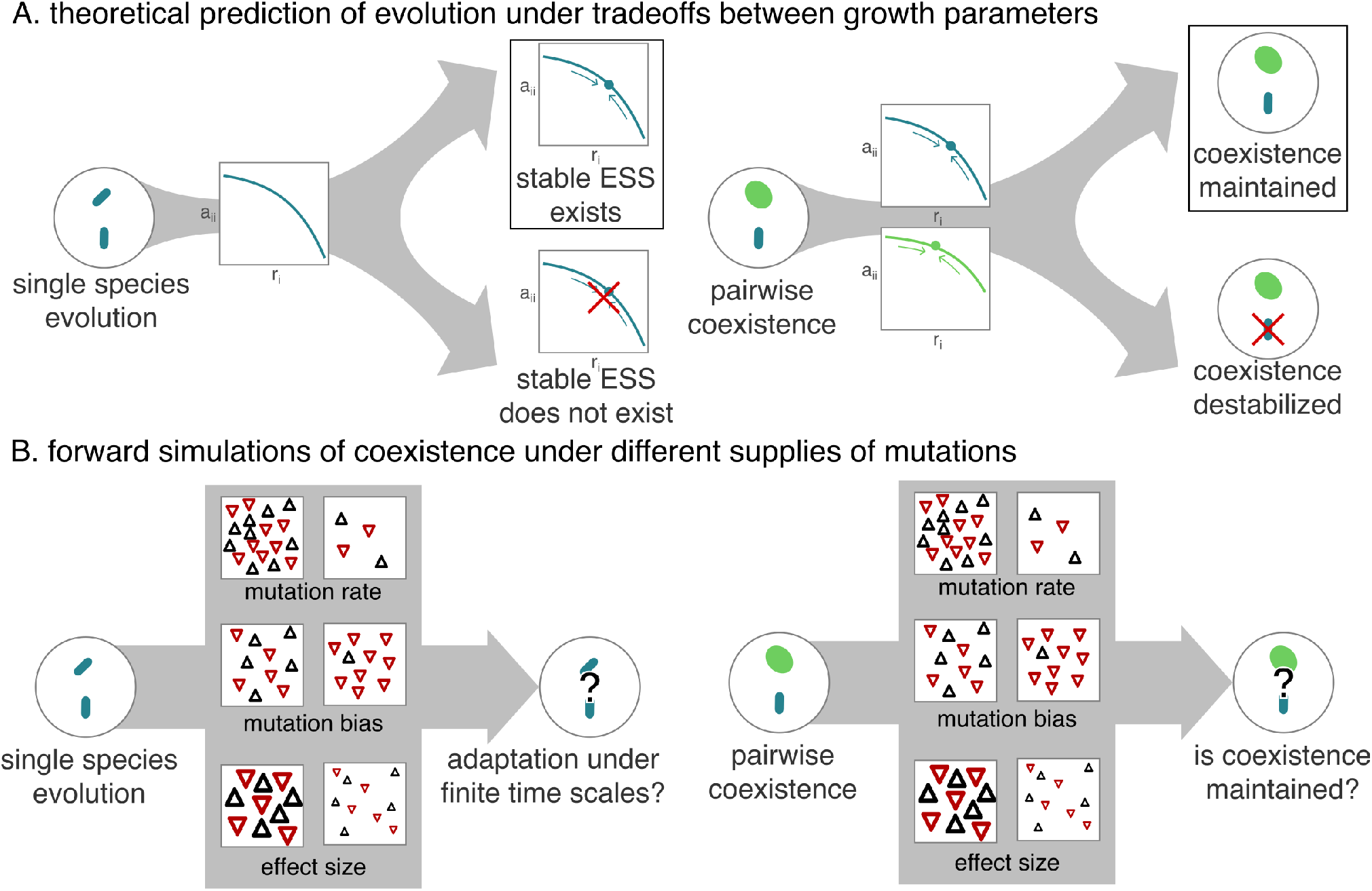
A. We use the adaptive dynamics framework to make theoretical predictions of evolutionary trajectory under trade-offs between growth parameters. B. We assess these predictions using forward simulations mapping the trajectory of evolution on varying the supply of mutations (mutation rate, supply of trait-increasing mutations (mutation bias), and mutation effect size), and calculate new quantities not available in adaptive dynamics, such as time-dependent fixation rates and the time to reach an Evolutionary Stable Strategy (ESS).

## Methods

### Model description and analysis

We used the adaptive dynamics framework to examine the fate of a mutation arising in a clonal population (Van Cleve, 2023). The growth equation that we used for both the single-species and paired-species cases was a form of the gLV equation (Mühlbauer et al., 2020; Lele et al., 2026), given as follows.

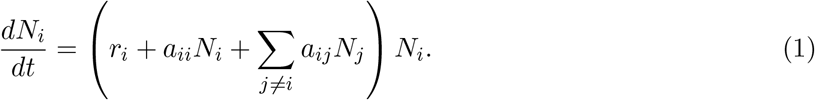

In this equation, *N*_*i*_ represents the abundance of species *i, r*_*i*_ is the basal growth rate, and *a*_*ii*_ and *a*_*ij*_ are coefficients that encompass all interactions each member of the focal species *i* can have. *a*_*ii*_ is the coefficient for intraspecific interaction (interaction of species *i* with itself), and *a*_*ij*_ are the coefficients for interspecific interaction (interaction of species *i* with all other species *j*). In evolutionary analyses, we supposed that *r*_*i*_ and *a*_*ii*_ values could evolve, while *a*_*ij*_ are fixed. This type of simple model is analytically tractable and reflective of the broadly studied *r*-*K* trade-offs in population ecology (Marshall et al., 2023). Models including evolution of *a*_*ij*_ values could be interesting candidates for future study (also see Bowers and White, 2002).

#### Single-species analysis

For the single species case (*i* = 1), we obtained the growth equations for the resident and an invading mutant,

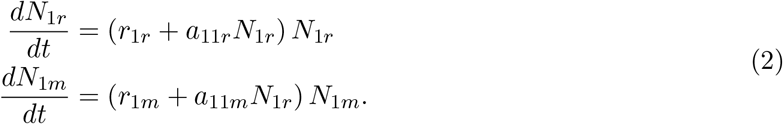

Here, the subscripts _*r*_ and _*m*_ denote resident and mutant parameters, respectively. We followed the standard adaptive dynamics approach of computing eigenvalues of the stability matrix and substituting the equilibrium value for the resident *N*_1*r*_ = − *r*_1_*/a*_11_ to determine the invasion fitness *s*_1*m*_ of the mutant. We find

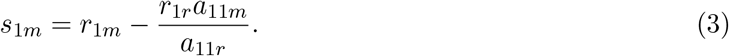

We examined the derivative of the invasion fitness *δs*^1*m*^*/δr*_1*m*_ to determine evolutionary outcomes, such as the existence of a candidate Evolutionary Stable Strategy (ESS), following standard adaptive dynamics approaches (Otto and Day, 2007; Brännström et al., 2013). We considered cases both with and without a trade-off function. For cases with a trade-off function (*a*_11_ = *f* (*r*_1_)), we additionally evaluated the nature of the candidate ESS using second-order partial derivatives, following standard adaptive dynamics methods (Bowers and White, 2002; Diekmann, 2002). We used pairwise invasibility plots (PIPs) to graphically check the analytical results (Supplementary Figure 1). We constrained our simulations to cases where a candidate ESS was predicted to exist. See Appendix section 3 for detailed derivations.

#### Multi-species analysis

We used a similar approach to obtain a candidate ESS both with and without a trade-off function for the two-species case. The system of growth equations included equations for both species, as well as a mutant arising in one species (here, assumed to be species 1) at a time, is given by

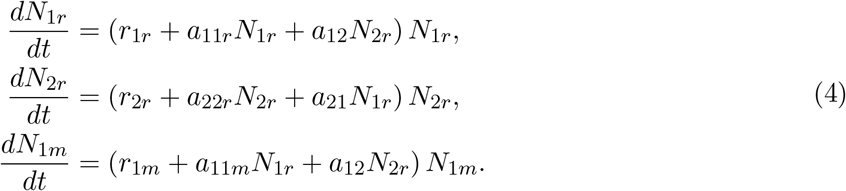

We obtained the invasion fitness from the eigenvalue of the stability matrix that contained the mutant parameters (Appendix A). We substituted the equilibrium values of the resident species *N*_1_ and *N*_2_ in the invasion fitness. Similarly, we obtained a symmetrical equation for invasion fitness for a mutant in species 2. The invasion fitness equations for both species are given by

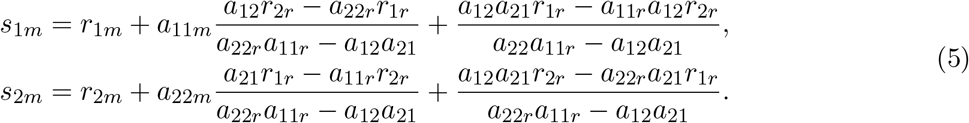

For the two-species case, we also imposed the additional condition that populations would be predicted to coexist at the initial condition. Without this condition, co-evolution would be unlikely to occur, as one species would be quickly extirpated even when both populations are initiated at positive population density.

Like the single-species case, we examined the partial derivative of the invasion fitness equations *δs*^1*m*^*/δr*_1*m*_ and *δs*^2*m*^*/δr*_2*m*_, using standard adaptive dynamics approaches to determine whether a candidate ESS would exist. To impose the condition that species must coexist during evolution, we used an established theoretical method to generate predictions of stable coexistence for pairs of species (Letten et al., 2017) that we adapted to our system of gLV equations in prior work (Lele et al., 2026). Once again, we considered cases with and without trade-off functions (*a*_11_ = *f* (*r*_1_) and *a*_22_ = *g*(*r*_2_)). Like before, we evaluated the nature of the candidate ESS using second-order partial derivatives, and used pairwise invasibility plots (PIPs) to graphically check the analytical results (Supplementary Figure 1). We constrained further analysis and simulations to trade-off functions where a) a candidate ESS was predicted to exist and b) species were predicted to coexist at the candidate ESS.

We used *Mathematica* v14.2 to evaluate our derivations above (Wolfram Research, Inc.). In cases where we were unable to solve equations analytically, we wrote Python scripts (v3.11.4) to solve them numerically for specific values of the constants. A detailed description of these analytical and numerical analyses can be found in the Appendix section 4.

### Simulation methods

We performed stochastic population-genetic simulations of species competing with gLV dynamics. We used these simulations to assess the extent to which deterministic predictions from adaptive dynamics would mirror results from simulations in which some model assumptions deviated from the assumptions of adaptive dynamics. For all simulations, we used `SLiM v4.3` (Haller and Messer, 2023) on the Tufts High-Performance Computation Cluster. All simulation code is available on GitHub and will be deposited into a stable repository at the time of publication.

#### Simulation model overview

The stochastic simulations include several features that differ from the mathematical assumptions of our adaptive dynamics calculations, and therefore could cause simulated outcomes to diverge from theoretically predicted outcomes. These include several standard assumptions within population genetics, specifically a) finite supply of mutations, b) finite population sizes, c) a specified distribution of effect mutations, which may include both small-effect and large-effect mutations, and d) the possibility for multiple effect mutations to segregate simultaneously. We made one additional assumption that is motivated by prior biological research into constraints on traits (Shoval et al., 2012; Pallasdies et al., 2021) – specifically, that trade-off functions represent upper bounds on performance that is determined by trait values. As such, trait values can fall off the trade-off curve, so long as fitness is bounded above by the trade-off curve (Sheftel et al., 2018). This view treats trade-off functions as limits on adaptation that are imposed by external, physical factors, and as such, the shape of the trade-off curve is best thought of as an output that depends both on the organism and the environment. We do not include pleiotropic constraints (*i*.*e*., anti-correlations between mutational effects on traits), but we consider possible future studies of pleiotropic models in the Discussion.

#### Trait model and mutation parameters

We used a non-Wright-Fisher model with one or two species and density-dependent reproduction. We used the same general framework for single-species and two-species simulations. All species in our simulations were haploid and recombination did not occur. Mutations arose independently in *r*_*i*_ and *a*_*ii*_ (*i* ∈ {1, 2}) (we consider pleiotropy between *r* and *a* in the discussion as a possible future direction, but do not include them here). We sampled effect sizes for *r*_*i*_ and *a*_*ii*_ from gamma distributions with mean 1 and shape parameter *α* = 0.5. We sampled trait-increasing and trait-decreasing mutations from separate gamma distributions, and the relative supply of either was determined by the parameter *m*_*k*_. Each effect allele had an additive impact on the growth rate and density-dependence of reproduction for each individual. Therefore, selection acts on trait values, and not directly on specific alleles.

We varied several factors in our analyses, including mutation rates, the magnitude of effect sizes, and the proportion of alleles that were trait-increasing or trait-decreasing (mutation bias). Mutation rates and mutation bias were controlled through default options in `SLiM`. We controlled the distribution of effect sizes through scaling factors on *r*_*i*_ and *a*_*ii*_, which we term *β*_*r*_ and *β*_*a*_. At any given time-point in the simulation, multiple values of *r*_*i*_ and *a*_*ii*_ may be represented in the population – for analysis, we tracked changes in the mean parameter values within each population to represent the consensus at a given time point.

#### Density-dependent growth

During each simulated generation, we calculated the parameter values for each individual by adding the quantitative effect of mutations (both trait-increasing and trait-decreasing) in the genome of each individual (scaled by the effect size) to the initial parameter value (which was constant as we assumed populations to be clonal initially) for *r*_*i*_ and *a*_*ii*_. Then, we used these updated parameter values to calculate the density-dependent per capita growth rate for each individual *k, z*_*k*_ = *dN*_*i,k*_*/N*_*i*_*dt*. We then take *z*_*k*_ + 1 as the mean in a Poisson distribution to determine the expected number of offspring for each individual. This allowed individuals with a higher per capita growth rate to potentially have more offspring and beneficial mutations to spread. We also included non-overlapping generations (to ignore potential complications with age-related effects), where the parents died at the end of each tick. If the per capita growth rate for all individuals was 0, the population would essentially replace itself and be stable. For all values of per capita growth rate below −1, no offspring were generated for that individual.

#### Trade-off functions

We incorporated trade-offs between *r*_*i*_ and *a*_*ii*_ by allowing mutations to accumulate independently in *r*_*i*_ and *a*_*ii*_, and used the trade-off function as an upper bound for the parameter values. Specifically, populations were allowed to accumulate mutations while the cumulative effect of mutations was bounded above by the trade-off curve. Once an individual adapted to reach the trade-offs, the effects of mutations on the trait values were constrained – individuals who accrued mutations that would result in trait values that exceeded the trade-off function were assigned trait values that lay on the trade-off function. This represents our interpretation of the trade-off function as a bound on performance that is governed by the environment, and hence no mutations can accrue to increase performance beyond these bounds. Populations were initiated with values of *r*_*i*_ and *a*_*ii*_ that fell either on or below the trade-off function (in the sense that fitness was bounded above by the trade-off curve), and we tracked the trajectories of these parameter values throughout the simulations.

The trade-off functions that we used were of the form 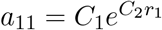 and 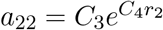. We considered multiple types of trade-off functions, including simple linear trade-offs, but opted for this specific mathematical form because it can result in an ESS for this specific type of model (see Appendix and Supplementary Figure 2). For our simulations, we chose values of *C*_1_, *C*_2_, *C*_3_ and *C*_4_ such that both species would coexist at the predicted single-species ESS as well as the two-species ESS, and the parameter values *r*_*i*_ and *a*_*ii*_ at the ESS were within the range of values sampled from sourdough microbes in previous work (Lele et al., 2026). Since microbial population density at equilibrium is typically very high (*>* 10^7^) and it is computationally infeasible to simulate such large populations, we rescaled our simulations. This required us to choose values of *C*_1_ and *C*_3_ such that the *a*_*ii*_ values at ESS were higher by a factor of 10^3^ compared to our measured values. This resulted in lower population sizes in the thousands while maintaining similar growth dynamics observed in empirical microbial interactions (Lele et al., 2026).

#### Selecting initial trait values and parameters

In all simulations, initial values for *r*_*i*_ and *a*_*ii*_ were chosen such that populations started away from their predicted ESS (that is, we assumed that they were initially maladapted). Additionally, the initial values for *r*_*i*_ were within the range of values sampled from sourdough microbes in our previous work as described above, while the values of *a*_*ii*_ were scaled by a factor of 10^3^ to maintain lower population sizes for computational feasibility (Supplementary figure 3). In general, we do not have empirical estimates of the number of sites (genome-wide) at which mutations can affect growth rates or competition coefficients in microbes, or the full distribution of effect sizes of mutations occurring at such sites. Consequently, we varied parameter values across a wide range of mutation rates, effect sizes, and the proportion of trait-increasing or trait-decreasing alleles to capture a large set of biologically plausible situations in our initial simulations. In subsequent simulations, we focused on a specific set of parameters, or smaller ranges of parameters, which we describe in more detail below. We tracked the population averages of trait values (*r*_*i*_ and *a*_*ii*_) and population size at fixed intervals throughout each simulation. Additionally, at the end of each simulation run we stored information about the mutations that were fixed in each population during the simulation, including the tick at which mutations arose, the time taken for a given mutation to fix, and the effect size of each fixed mutation.

**Table 1:**
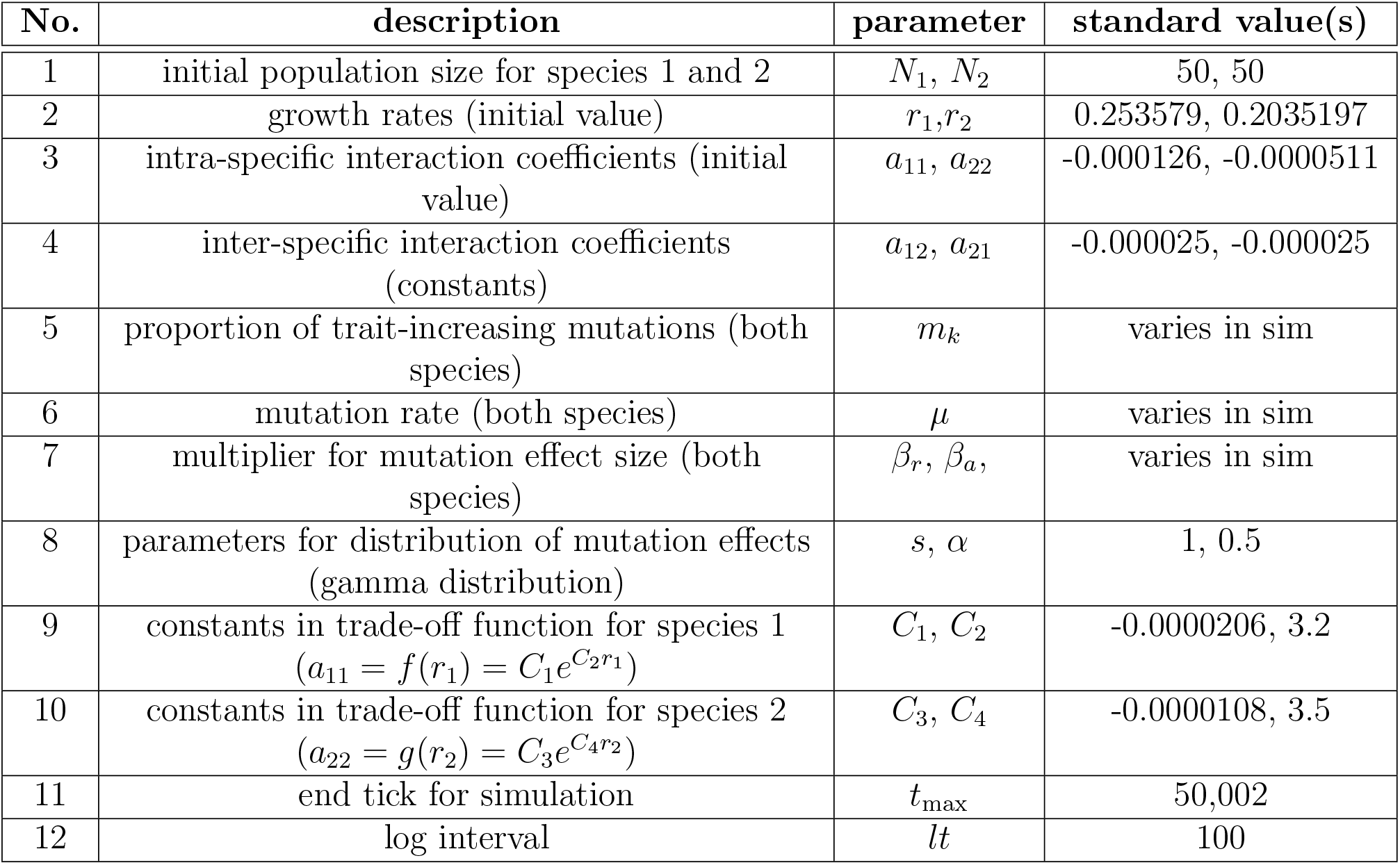
Table containing standard parameters for simulations where one or two species evolve under given trade-off functions between growth parameters *r*_1_ and *a*_11_, and *r*_2_ and *a*_22_ (Figures 3 and 4).

#### Finite-time simulations (both single and two species)

For the first set of simulations, we used a single pair of initial values for *r*_*i*_ and *a*_*ii*_. This allowed us to investigate the impacts of other parameters (such as mutation rates and effect sizes) on the outcome independent of initial conditions, which we vary in subsequent analyses. In these simulations, we varied the mutation rate (*µ*) between 10^−8^ to 10^−6^, the proportion of trait-increasing mutations (*m*_*k*_) from ^1^*/*2 to ^1^*/*500, and the scalar multiplier on effect sizes from 0.1 to 0.005 for *r*_*i*_ (*β*_*r*_) and 10^−5^ to 5×10^−7^ for *a*_*ii*_ (*β*_*a*_). We ran 100 simulations for each parameter combination for 5 × 10^4^ “ticks” (`SLiM` nomenclature for generations), and tracked mean trait values and population size at intervals of 100 ticks throughout the simulation (see Table 1 for these parameter values and Table 1 in Supplementary information for additional information). We conducted two kinds of simulations - single-species simulations, and two-species simulations, where we extended our single-species simulations to a pair of interacting species. The results of these simulations are discussed in the Results sections “Rate of adaptation depends on mutation rate, effect size and supply of trait-increasing mutations in populations of single species” and “Expected coexistence patterns are often in conflict with stochastic simulations”, and Figures 3 and 4.

For the next set of simulations, we varied the initial values of *r*_*i*_ and *a*_*ii*_ between simulations, and kept all other parameters consistent (see supplementary Table 2 for parameter values) between simulation runs to assess how initial condition choices might affect our analyses. This allowed us to evaluate how initial parameter values influenced the destabilization of coexistence. We picked initial values of *r*_*i*_ and *a*_*ii*_ that led to coexistence, and added an additional constraint that the scaled Euclidean distance between the initial parameter values and the ESS parameter values must be between 1 and 5, to allow the opportunity for substantial adaptation. We calculated scaled distances with the following formula, 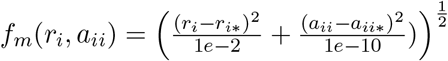, which treats changes in *r*_*i*_ and *a*_*ii*_ equally since these parameters differ substantially in magnitude. We ran 100 simulations for each value of *r*_*i*_ and *a*_*ii*_, and tracked the population size and the population mean values of *r*_*i*_ and *a*_*ii*_ at intervals of 100 ticks throughout the simulation. The results from these simulations are discussed in Results section “Initial growth parameters influence the probability that coexistence is destabilized”, and Figure 5.

#### Conditioning simulation end-points on pre-selected criteria

Most simulations in this study were terminated at a fixed endpoint (typically 5 × 10^4^ generations), which we took as representative of a long term evolution study (Lenski, 2023). We also sought to perform simulations in which the termination conditions corresponded to reaching the ESS or extinction – conditioning on these criteria allows us to compare to adaptive dynamics more directly, since it effectively removes the possibility of non-convergence to the ESS in finite time.

We terminated these simulations soon after the ‘end conditions’ of adaptation were reached, which we define here as either when one species went extinct, or the distance between population mean values and ESS values of *r*_*i*_ and *a*_*ii*_ was less than 0.5. To calculate this distance, we used the same distance function *f*_*m*_(*r*_*i*_, *a*_*ii*_) described in the prior section. When end conditions were reached, the simulation continued for 1,000 additional ticks and was then terminated. If the end conditions were never reached, simulations terminated after 500,000 ticks.

For this set of simulations, we varied the mutation rate uniformly between 10^−7^ to 10^−6^, and kept all other parameters consistent between simulations (see Supplementary Table 3 for parameter values). Mutation rates varied independently for both species. We ran two sets of simulations, one in which the proportion of trait-increasing mutations was ^1^*/*2, and one in which the proportion of trait-increasing mutations was ^1^*/*10. As before, we tracked the population size and the population mean values of *r*_*i*_ and *a*_*ii*_ at intervals of 100 ticks throughout the simulation. We ran 5 × 10^4^ simulations for both proportions of trait-increasing mutations. For simulations where the proportion of trait-increasing mutations was ^1^*/*2, 15 simulations did not reach end conditions (0.03%). For simulations where the proportion of trait-increasing mutations was ^1^*/*10, 902 simulations did not reach end conditions (1.8%). We excluded all simulations that did not reach end conditions in subsequent analyses. The results from these simulations are discussed in Results section “Asymmetry in the mutation spectrum influences coexistence outcomes”, and Figure 6.

## Results

### trade-off functions predict stable ESS in adaptive dynamics analyses

We used the adaptive dynamics framework to predict the evolutionary trajectory of a clonal population consisting of a single species evolving under gLV dynamics. Our goal in this analysis was to establish conditions under which evolution would proceed to an ESS, which we would then use as a theoretical baseline to which we can compare population-genetic simulations. As expected, when *r*_1_ and *a*_11_ evolved without constraints, populations were predicted to evolve towards higher values of *r*_1_, and less negative values of *a*_11_ in an unbounded manner (see Appendix and Supplementary Figure 1).

We then constrained our analysis by imposing trade-off functions *a*_11_ = *f* (*r*_1_) that can generate a candidate ESS. We considered a range of shapes for trade-off functions, before choosing a trade-off function of the form 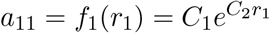 (Supplementary Figure 2) since it led to a candidate ESS that was convergence and evolutionarily stable at *r*_1_ = ^1^*/C*_2_ (Figure 2A, B). By examining the invasion fitness landscape and performing additional calculations, we confirm that this ESS is convergence and evolutionarily stable, indicating that populations will evolve to the ESS when the assumptions of adaptive dynamics are met.

**Figure 2:**
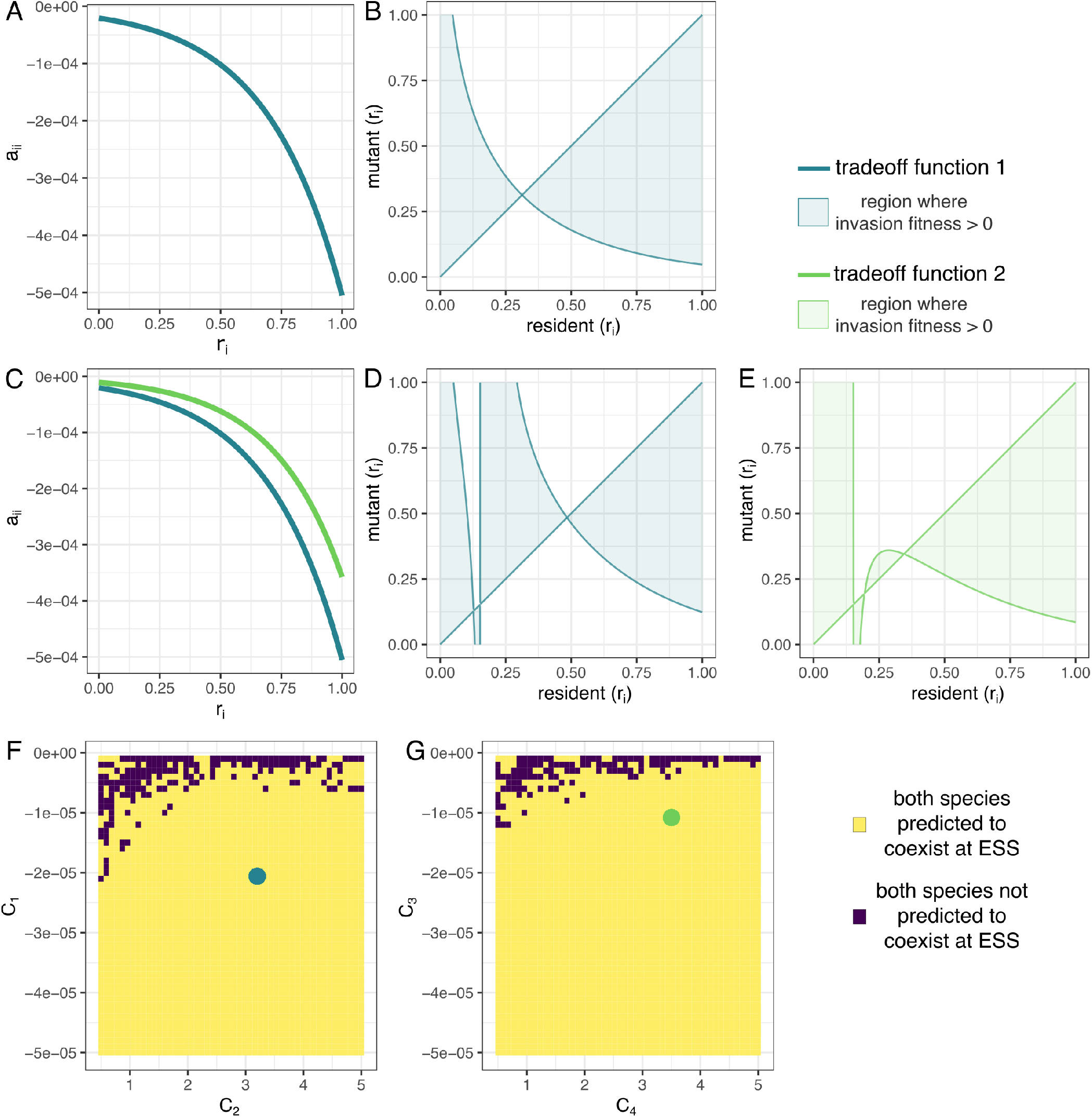
Adaptive dynamics analysis with trade-offs between *r*_*i*_ and *a*_*ii*_. A. Trade-off function 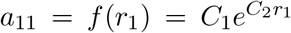. B. Pairwise invasibility plot for *r*_1_ under the Trade-off function in A, showing the existence of a candidate ESS at *r*_1_ = 0.3125. C. Trade-off functions for pairs of species, species 1: 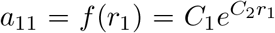, species 2: 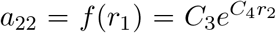. D. Pairwise invasibility plot for species 1, *r*_1_ under the trade-off function in C, showing the existence of a candidate ESS at *r*_1_ = 0.498344. E. Pairwise invasibility plot for species 2, *r*_2_ under the trade-off function in C, showing the existence of a candidate ESS at *r*_2_ = 0.362690. F and G. Predictions of coexistence for two species evolving under trade-off functions of the form 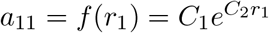 and 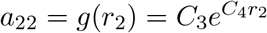. Values of *C*_1_, *C*_2_, *C*_3_, and *C*_4_ are given in Table 1.

Using the same approach, we obtained predictions for the evolutionary trajectory of a community consisting of clonal populations of two species. Similar to the single species case, we found that when mutations accumulated independently in *r*_*i*_ and *a*_*ii*_ for both species, populations were predicted to evolve towards higher values of *r*_*i*_, and less negative values of *a*_*ii*_ in an unbounded manner (see Appendix and Supplementary Figure 1). When we introduced trade-off functions of the form 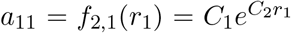 and 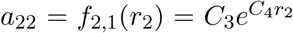, we obtained a candidate ESS (Figure 2C-E). From the pairwise invasibility plot and additional calculations, we confirm that this candidate ESS is both convergence stable and evolutionarily stable. Additionally, we confirmed that both species were predicted to coexist at the ESS for a range of values of the constants *C*_1_, *C*_2_, *C*_3_, and *C*_4_ (Figure 2F-G), from which we picked specific values that we used for all simulations below (Table 1).

In sum, unconstrained evolution results in run-away adaptation under the generalized LV model, while there exist trade-off functions that result in an ESS in both the single-species and two-species cases, as expected.

### Rate of adaptation depends on mutation rate, effect size and supply of trait-increasing mutations in populations of single species

#### Populations initialized on the trade-off curve (Population genetics theory)

As a first step in linking the adaptive dynamics framework to a mechanistic population genetic model, we investigated the rate of change in both trait space and sequence space for a single species evolving under constrained LV dynamics. Standard adaptive dynamics theory provides insight into the existence of the ESS, but the rate of evolution additionally depends on the effect sizes and mutation rates of effect alleles for the focal traits, *r* and *a*, which can be calculated using population genetics theory. For simplicity, we perform this calculation from the perspective of mutations in growth rate *r*, noting that this choice has no substantial effect on the outcome because *a* is determined by *r* when evolution occurs along the trade-off curve and hence there is only one degree of freedom.

To calculate the time required to transit between an initial condition *r*_*i*_ to a final condition *r*_*f*_ along the trade-off curve, we supposed that the population was initialized away from the ESS. We suppose in these calculations that *r*_*i*_ is lower than the growth rate at the ESS, but similar logic can be applied to initial conditions in which *r*_*i*_ *> r*_ESS_. As in standard population genetics theory, the rate of adaptation therefore depends on the strength of selection acting on new mutations (via their effect sizes on *r* and *a*) and the supply of mutations. However, population sizes are also contingent on the evolutionary dynamics, since the equilibrium population size is given by *N*_eq_ = ^*r*^*/a*, resulting in coupled ecological and evolutionary dynamics. Our calculation proceeds by supposing that the time to reach the ESS can be approximated by summing the cumulative effects of mutations on one of the focal traits, and that populations equilibrate to their expected size rapidly.

Following standard population genetics theory, we find that the transit time from an initial value of *r*_*i*_ to a final value *r*_*f*_ is given by

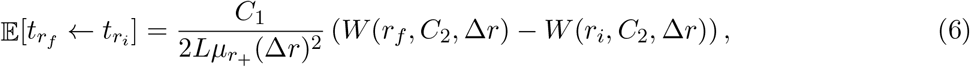

where *W* (*r, C*_2_, Δ*r*) is a complex function that depends on *r*, the effect size Δ*r*, and the trade-off function via *C*_2_. The form of *W* is given in the Appendix. We validated our calculation by comparing expected values of 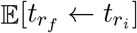 to values obtained from stochastic simulation, as well as time-dependent fixation rates (Fig. S4).

Examining eqn. 6, we observe that the time to reach the ESS is inversely proportional to the genomewide mutation rate for *r*-increasing mutations 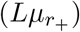 and the square of the effect size of mutations (Δ*r*)^2^. This is expected – higher rates of mutation will decrease the time to reach the ESS, while larger effect sizes will increase the rate both through increased probability of fixation and larger step sizes in trait space per fixation event. This suggests that changes in the distribution of effect sizes may have a larger impact on the rate of adaptation than changes in the size of the mutational target. However, this scaling on effect sizes is somewhat more complex given that *W* (*r, C*_2_, Δ*r*) also depends on Δ*r*. We show in the Appendix that *W* (*r, C*_2_, Δ*r*) only varies modestly as a function of effect size Δ*r*, and consequently it has a smaller effect on the transit time than the (Δ*r*)^2^ term in the denominator of eqn. 6. Larger values of the trade-off coefficient *C*_1_ will increase the time to reach the ESS. An obvious consequence of this time-dependence is that many populations evolving along the trade-off curve may fail to reach their predicted ESS in finite time. For a population with 2*Nµ*_+_ ≈ 2.5*e* − 5 and *L* = 10^3^, our calculations suggest that it can take in excess of 10^4^ generations to reach the ESS, depending on the magnitude of mutational effects on *r*. (Supplementary Figure 4).

#### Populations initialized off the trade-off curve (Population genetics simulations)

Having obtained an expression for the time to reach the ESS for populations evolving *along* the trade-off function, we next explored the transit time to the ESS for populations initiated at parameter values that fall *off* the trade-off function. Analytical calculations are more challenging for populations that are initiated off the trade-off curve, because of the multitude of paths that can be taken to reach the ESS, in contrast to populations that evolve along the trade-off curve. We performed finite-time simulations in `SLiM` for these analyses, varying the parameters that determined the effect size and occurrence of mutations (Figure 3A).

We calculated the time at which population trait values arrived at the ESS in each simulation replicate (see Methods; Figure 3A). As expected (and similar to our findings in the prior section), lower mutation rates, lower supplies of trait-increasing mutations, and smaller effect sizes all caused populations to fail to reach the predicted ESS in the finite time scales of our simulations. We also calculated the minimum distance between the population mean growth parameters and the ESS parameters at the end of each simulation, and found concordant patterns (Supplementary Figure 5). These findings are intuitive – however, we observed large variance across replicates in the time taken to reach ESS for intermediate parameter values, suggesting that experimental replicates could be highly variable despite evolving on the same selection surface. This is evident in the trajectories of the population mean values of *r*_*i*_ and *a*_*ii*_ (Figure 3B-C). For simulation runs in which most populations reached the ESS, we observed that among the mutations that reach fixation in *r*_*i*_ and *a*_*ii*_, larger effect trait-increasing mutations drive adaptation to the ESS earlier, while the mutations that fix later are of smaller effect sizes, especially for mutations in *r*_*i*_ (Figure 3B, Supplementary Figure 5). This is consistent with our theoretical calculations in the prior section, since only very small effect mutations are beneficial when trait values are close to the ESS. Effect sizes have the largest impact on the time time taken to reach the ESS – among the simulations with the smallest effect sizes, only simulations with the largest proportion of trait increasing mutations had any fixation events. For simulations with larger effect sizes, we observe a modest impact of mutation rates on the time taken to reach the ESS.

**Figure 3:**
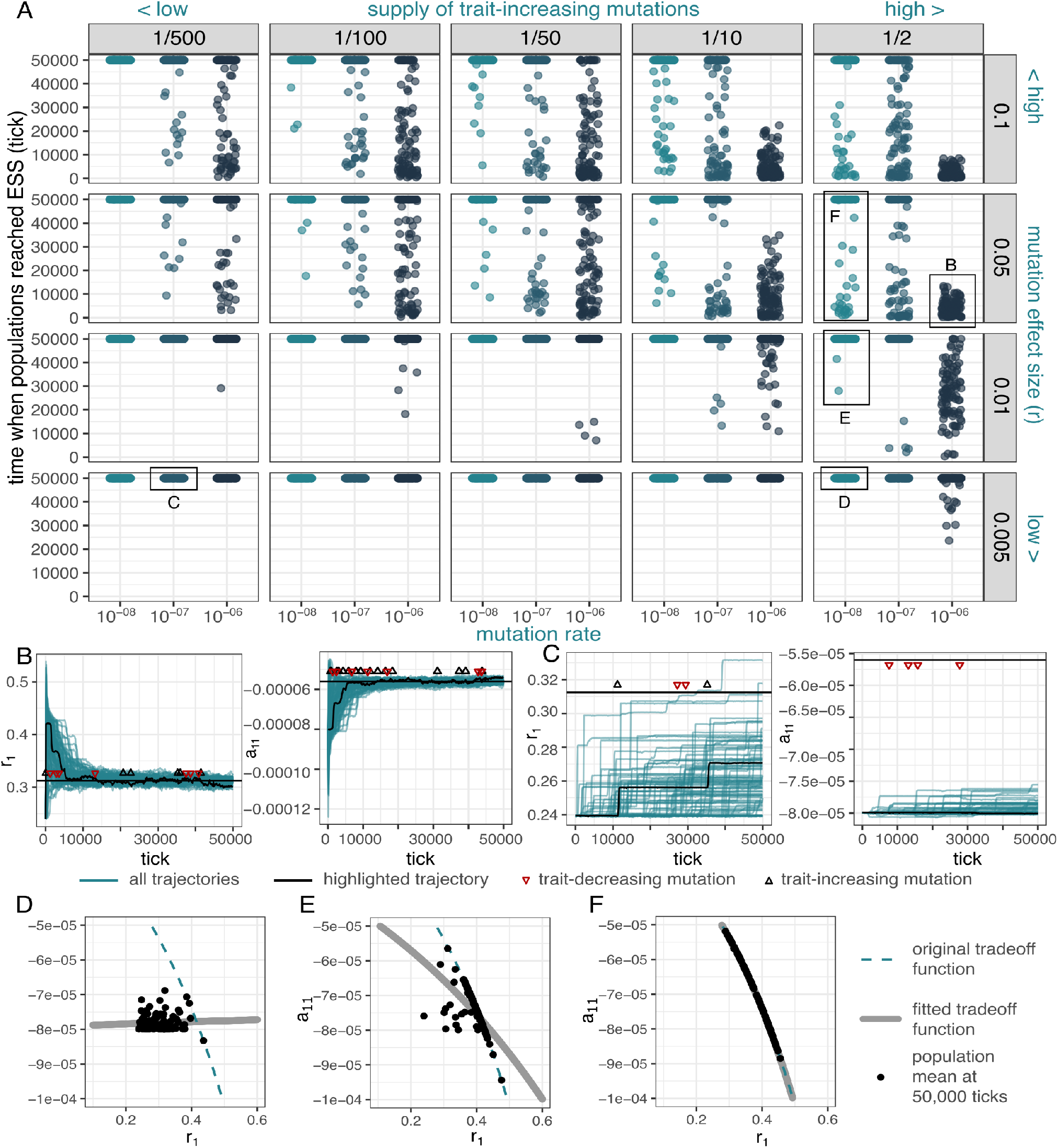
Single-species evolution over 50,000 generations. A. Time at which individual simulation runs (*n* = 100) reached the ESS for various values of *µ, m*_*k*_, and *β*_*r*_. Colors indicate different mutation rates (also on x-axis). Values of 50,000 indicate populations that never reached the ESS. B. Population mean values of *r*_1_ and *a*_11_ over time (*µ* = 10^−6^, *m*_*k*_ = ^1^*/*2, *β*_*r*_ = 5*e* − 2, *β*_*a*_ = 5*e* − 6). C. Population mean values of *r*_1_ and *a*_11_ (*µ* = 10^−7^, *m*_*k*_ = ^1^*/*500, *β*_*r*_ = 5*e* − 3 and *β*_*a*_ = 5*e* − 7). In B and C, colored lines indicate trajectories for individual replicates. Black lines are an example replicate corresponding to the plotted mutation events. Red (black) triangles indicate fixations of trait-decreasing (trait-increasing) alleles. D,E,F. Estimating trade-off functions from simulated data with *m*_*k*_ = ^1^*/*2, *µ* = 10*e*−8, and *β*_*r*_ = 5*e*−3, *β*_*a*_ = 5*e*−7 (D), *β*_*r*_ = 0.01, *β*_*a*_ = 1*e*−6 (E), *β*_*r*_ = 5*e* − 2, *β*_*a*_ = 5*e* − 6 (F). Black dots indicate population mean values of *r*_1_ and *a*_11_, blue dashed line is the true trade-off function, and gray lines are best-fit trade-offs.

#### Detection of growth/competition trade-offs also depends on supply of mutations in finite time scales

Given the multitude of paths that simulations took through (*r, a*) space, we also queried whether it would be possible to detect growth/competition trade-offs by comparing *r* and *a* values across replicates, an analysis similar to those performed in prior experimental evolution studies (Novak et al., 2006). We used the observed (*r, a*) coordinates at the simulation endpoint for each replicate evolved from a common initial condition as input data in an attempt to infer the trade-off function (Figure 3D-F, Supplementary Figure 6). For lower values of mutation rates, proportion of trait-increasing mutations, and mutation effect size, adaptation proceeded slowly enough that populations were far away from the trade-off curve at the endpoint of the simulation ticks (Figure 3D), and the estimated trade-off curve bore little resemblance to the true trade-off. However, when we increased the supply of mutations, we found that the mean values of growth parameters were bound by the trade-off function, and hence the shape of the estimated trade-off function more closely resembled the true trade-off function (Figure 3E-F). This implies that studies seeking to infer trade-offs from experimental evolution replicates could be severely limited in their ability to detect the curve, depending on the underlying evolutionary parameters, even after tens of thousands of generations.

### Expected coexistence patterns are often in conflict with stochastic simulations

Next, we investigated the impact of mutation spectra on coexistence outcomes for pairs of species. We hypothesized that some combinations of mutation rates and effect sizes would result in competitive exclusion, even when adaptive dynamics predicts pairwise coexistence. For example, this is likely to occur in the limit of very low mutation rate in one species, because its competitor may rapidly transit the parameter space while the focal species remains stationary. However, the precise conditions under which such dynamics will occur are non-obvious and depend on the stochastic trajectories through trait space. To investigate competition outcomes with finite populations, we extended the `SLiM` simulations we used in the previous section to accommodate pairs of species that accumulated mutations and evolved together, such that their growth was governed by gLV equations.

In finite time simulations, neither species went extinct for low mutation rates, low proportions of traitincreasing mutations, and smaller mutation effect sizes (Figure 4A, bottom left). At high mutation rates, proportions of trait-increasing mutations, and mutation effect sizes, once again neither species went extinct for the duration of the simulations (Figure 4A, top right). However, we observed an increase in replicates in which one species went extinct for intermediate values of the mutation supply parameters (Figure 4A). Extinctions also occurred earlier in the simulations for these intermediate parameter values.

**Figure 4:**
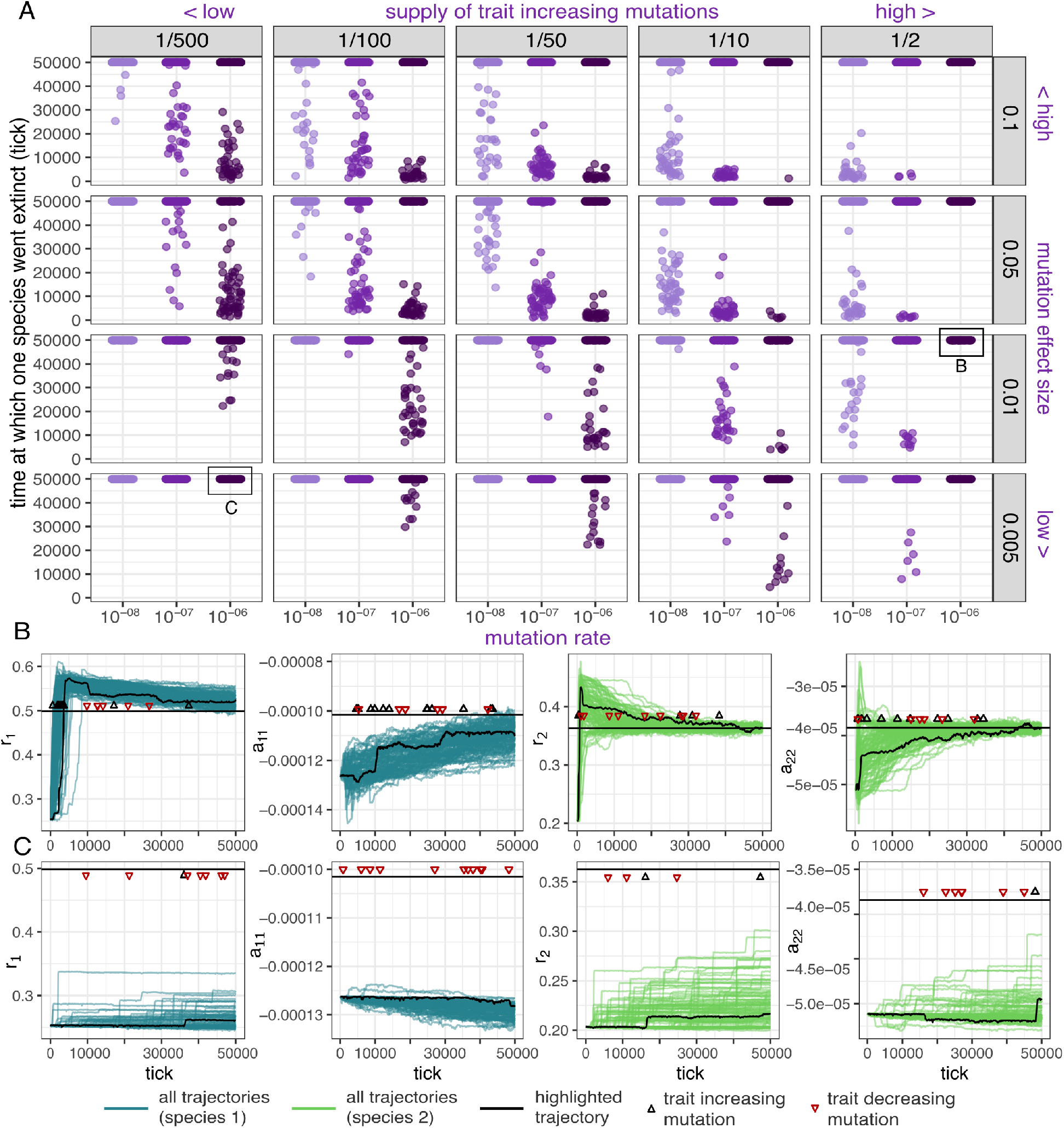
Two-species coevolution over 50,000 generations. A. Time at which one population went extinct in individual simulation runs (n=100) for different mutation rates (*µ*), proportion of beneficial mutations (*m*_*k*_), and mutation effect sizes (*β*_*r*_ and *β*_*a*_). Colors indicate different mutation rates (also on x axis). Populations at 50,000 did not go extinct before the end of the simulation. B. *r*_1_, *a*_11_, *r*_2_ and *a*_22_ over time for simulations with *µ* = 1*e* − 6, *m*_*k*_ = ^1^*/*2, *β*_*r*_ = 1*e* − 2 and *β*_*a*_ = 1*e* − 6. C. *r*_1_, *a*_11_, *r*_2_ and *a*_22_ over time for simulations with *mut* = 1*e* − 6, *m*_*k*_ = ^1^*/*500, *β*_*r*_ = 5*e* − 3 and *β*_*a*_ = 5*e* − 7. Colored lines indicate trajectories for all simulations (blue for species 1, green for species 2), black lines indicate trajectories for one replicate for which mutations that occurred in the population are shown. Red triangles indicate a trait-decreasing mutation and black triangles indicate a trait-increasing mutation at the given tick.

The lack of extinctions at lower values of mutation supply parameters could be explained by a general failure to adapt substantially from the initial conditions for the duration of the simulation. Sometimes, this even resulted in maladaptation through the accumulation of deleterious alleles (Figure 4C, Supplementary Figure 7). Indeed, we observe that over much longer time scales (500,000 ticks), a subset of simulations in this category went extinct, highlighting that finite time scales were crucial to maintain coexistence (Supplementary Figure 7). Conversely, in the upper range of mutation supply parameters, both species had high adaptive capacities and evolved to the ESS rapidly, maintaining coexistence throughout the trajectory (Figure 4B). Like the single-species case, rapid adaptation was driven by the fixation of larger-effect trait-increasing mutations earlier in adaptation.

### Initial growth parameters influence the probability that coexistence is destabilized

We focused on a single set of initial parameter values (*r*_1_, *a*_11_, *r*_2_ and *a*_22_) in the prior section, but destabilization of coexistence could also be mediated by initial conditions. Such dependence would not be captured by adaptive dynamics, which does not specifically address initial conditions except to assume that the system is close enough to the ESS (or another fixed point) that the specific conditions can be ignored. However, if parameter values along the evolutionary trajectory fall outside the coexistence region, then one of the species might go extinct. Alternatively, the trajectories between initial parameters (assumed to generate coexistence) and the ESS parameter values might preclude this possibility or make it exceedingly unlikely. Such conditions may be especially likely to occur when one species has much lower adaptive capacity to the other.

We tested this possibility by randomly sampling a set of 100 initial parameter values for both species inside the region of coexistence. For all these values, we computed the niche overlap (*ρ*) and fitness difference (*f*_1_*/f*_2_) when both species were at the initial conditions, and at the candidate ESS supposing that only one of the two species was allowed to evolve. When niche overlap is high or when fitness differences are very small, coexistence may be prohibited. Prior research relates the parameters of gLV models (*r* and *a*_*ij*_) to niche and fitness overlap, allowing for simple predictions of coexistence in pairwise competitions (Chesson, 2000; Letten et al., 2017). The (*ρ,f*_1_*/f*_2_) coordinate at the ESS when only one species evolves corresponds to the case in which only species 2 has a high enough supply of beneficial mutations to evolve rapidly, while species 1 is stationary at its initial condition. We then ran simulations for these initial parameter values, picking values of mutation rate, supply of beneficial mutations, and mutation effect size that led to extinctions in the previous section (see Table 2 in Appendix). We observe that simulations in which the initial parameter values were predicted to lead to potential extinctions, were indeed more likely to result in extinctions (Figure 5). We also observed that simulations in which the ESS fell inside the coexistence region but was close to the boundary for competitive exclusion sometimes resulted in extinctions. Finally, we observed that due to our choice of trade-off functions, there was an asymmetry between the extinction risk for both species. This asymmetry was reflected in the predictions of coexistence when species 2 remained stationery at initial parameter values, while species 1 evolved rapidly (Supplementary Figure 8), which did not predict extinctions, and was reflected in our simulations, where species 1 consistently went extinct.

**Figure 5:**
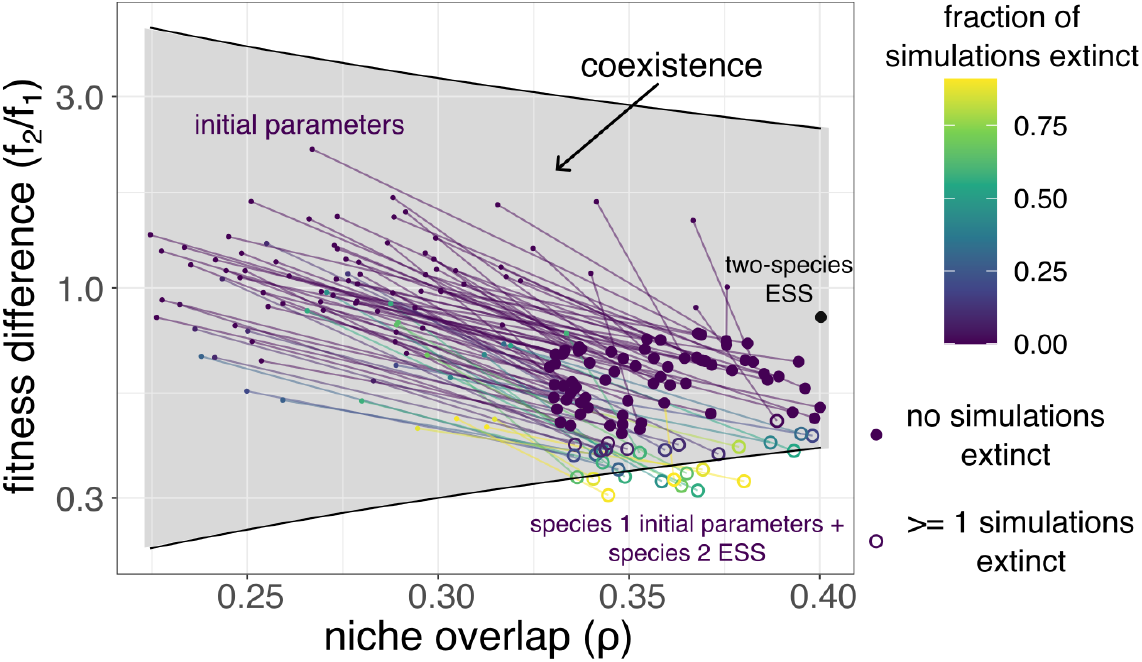
Predictions of coexistence between 100 randomly chosen initial parameter combinations of two species, along with predictions for species 1 initial parameters and species 2 ESS. Initial parameter values are shown with small circles, larger circles indicate ESS parameter values. Open circles indicate parameter combinations where at least one replicate resulted in extinction. The color of the open circle represents the fraction of simulations in which one species went extinct after 5 × 10^4^ generations.

### Asymmetry in the mutation spectrum influences coexistence outcomes

So far, we have largely focused on the outcomes of evolution within finite timescales, typically capped at 5 × 10^4^ generations (a timescale similar to the longest running microbial evolution experiments, Lenski (2023)). Though finite time simulations are relevant because all experiments are necessarily finite in duration, we sought to perform simulations of variable time in which termination is conditioned on either reaching the ESS or extinction, and each species’ mutation spectrum could vary independently. These simulations are important for assessing the relationship between population genetic models and adaptive dynamics, since predicted outcomes in adaptive dynamics are not time-dependent. In this set of simulations, we independently varied the mutation rate (*µ*) for both species, picking values from a uniform distribution between 10^−7^ and 10^−6^.

In general, we observed that more replicate simulations went extinct for lower values of *µ* across both proportions of trait-increasing mutations (Figure 6A,B). Additionally, overall extinction rates increased when the proportion of trait-increasing mutations decreased (Figure 6B). Results were asymmetric with respect to the two simulated species – the proportion of populations that went extinct were much higher when species one had a lower mutation rate compared to species two, while very few simulations resulted in extinctions when species one had a higher mutation rate than species two. This asymmetry could be due to the predicted coexistence between species 2 ESS and species 1 initial parameters being closer to the coexistence boundary (Figure 5, Supplementary Figure 8). Thus, when mutation rates were lower in species 1 and higher in species 2, a higher proportion of simulations ended due to species one going extinct. When mutation rates were higher for species 1 and lower for species 2, proportions of extinctions were very low.

**Figure 6:**
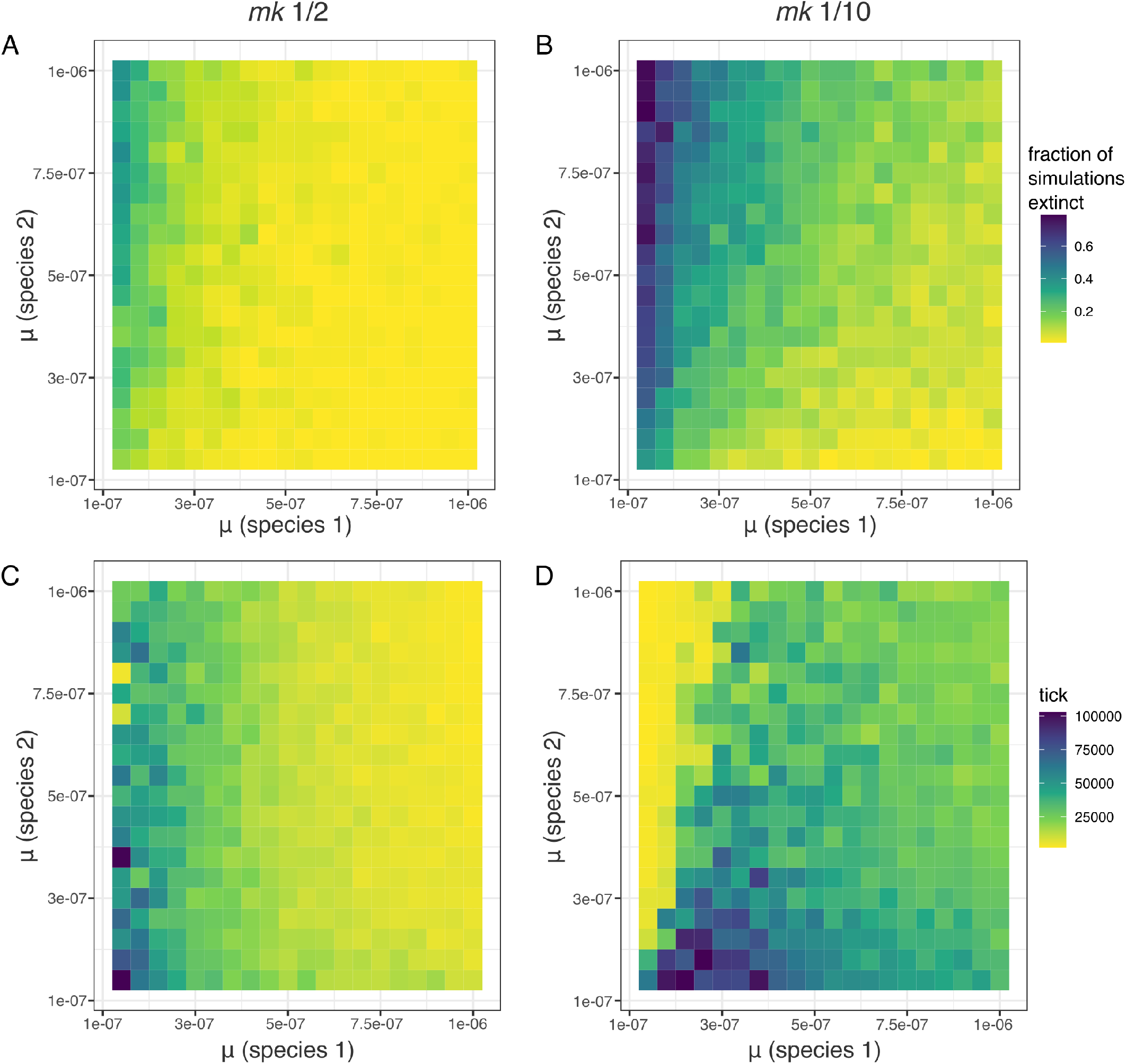
Two-species coevolution, conditioned on reaching the ESS or extinction. A. Fraction of simulations in which one species went extinct, for mutation rates varying over 10^−6^ to 10^−7^ and *m*_*k*_ = ^1^*/*2. B. Fraction of simulations in which one species went extinct, for mutation rates varying over 10^−6^ to 10^−7^ and *m*_*k*_ = ^1^*/*10. For A and B, color intensity indicates fraction of simulations in which populations eventually went extinct. C. Median tick at which replicate simulations reached end conditions, for mutation rates varying over 10^−6^ to 10^−7^ and *m*_*k*_ = ^1^*/*2. D. Median tick at which replicate simulations reached end conditions, for mutation rates varying over 10^−6^ to 10^−7^ and *m*_*k*_ = ^1^*/*10. For C and D, color intensity indicates median value.

Similar to the simulations in the earlier time scales that ran for a finite duration, replicates with lower mutation rates took much longer to reach the termination conditions (Figure 6 C,D). However, for the populations where a much larger fraction went extinct, we saw a dramatic decrease in the median time of simulation end, indicating that when extinctions happened, they happened earlier during the adaptation process. This pattern was especially prominent when the supply of beneficial mutations was much lower (Figure 6D).

## Discussion

While adaptive dynamics theory has provided major insights into evolution over invasion fitness landscapes, it has long been appreciated that its results might conflict with mechanistic models of evolutionary processes. Studies connecting population genetics to adaptive dynamics therefore have the potential to improve our fundamental understanding of evolution (and coevolution) by combining the strengths of both approaches – specifically, by leveraging an understanding of eco-evolutionary outcomes obtained with adaptive dynamics and quantitative predictions about commonly collected experimental/sequence data that are readily available from population genetics. In this study, we used a data-driven model of competition between microbes to show that population genetic models often recapture predictions made by adaptive dynamics, such as evolution towards an ESS. However, in finite time and finite populations there are parameter-dependent discrepancies between the modeling frameworks, several of which are relevant to the interpretation of experimental evolution studies. Some of our results are intuitive – specifically, the time to reach the ESS depends on the inverse of the square of the effect size of fitness-increasing mutations and the inverse of the rate of supply of such mutations, similar to established population genetic theory (Equation 6). Other results are potentially less obvious – for example, the expectations of coexistence or a stable ESS obtained with adaptive dynamics are often unrealized when there are large discrepancies in the supply of large-effect, trait-increasing mutations between species, but only if intermediate evolutionary trajectories are likely to take the system outside of the coexistence region (Figures 5-6).

Trade-offs between traits related to growth and competition may be major features of evolution along fitness landscapes (Stearns, 1998), and microbes are particularly well-suited to characterize these trade-offs experimentally. The idea that the evolution of traits that increase growth rates must be constrained by other costs makes intuitive sense and has strong theoretical support from both evolutionary (Stearns, 1998; Roff and Fairbairn, 2007) and metabolic models (Pfeiffer and Bonhoeffer, 2002; Frank, 2010). In fact, many experimental studies find that there are trade-offs between growth parameters (*r* − *K* trade-offs) (Jasmin and Zeyl, 2012; Maharjan et al., 2007; Bachmann et al., 2013; Meyer et al., 2015; Moerman et al., 2020). However, other experiments sometimes fail to find evidence of these trade-offs, instead observing positive correlations between *r* and *K* (Wei and Zhang, 2019; Reding-Roman et al., 2017; Marshall et al., 2022). This diversity of observed *r* − *K* trade-offs might reflect different biological mechanisms or environmental conditions might be at work in these situations, better captured through *r* − *a* relationships (Marshall et al., 2023). Yet, as our modeling shows, the failure to detect trade-offs could also simply reflect a low supply of trait-altering mutations (Figure 3). While it is clear in the limit of very low mutation rates that trade-offs will not be detected because species will evolve slowly, our study shows that this can occur even on very long experimental timescales with thousands of generations and data-driven parameters for the underlying species interactions.

Previous research in adaptive dynamics has also shown that trade-off functions govern the trajectory and outcome of adaptation (Kisdi, 2006, 2015). In this work, we found that the existence and nature of the ESS depends on the choice of trade-off function. We chose to focus on an exponentially decreasing trade-off between *r*_*i*_ and *a*_*ii*_, which leads to a trade-off between *r* and *K* that is positive at low growth rates and negative at high growth rates. This complex relationship could exist due to the different strategies species can utilize at the ends of the growth rate spectrum (Lipson, 2015), and allows the possibility of both positive and negative growth-yield trade-offs to be detected under different evolutionary histories (Novak et al., 2006). However, other *r*_*i*_ − *a*_*ii*_ trade-offs are possible, and another potential trade-off that we did not investigate further also led to a stable ESS, at least in the single-species case (Supplementary Figure 1). Additionally, it would be interesting to further investigate how populations would evolve under trade-offs that did not predict a stable ESS.

Our population genetic simulations differed from the assumptions we made under the adaptive dynamics framework in several important ways, sometimes resulting in discrepancies between predicted and observed outcomes. First, we assumed that the trade-off between *r*_*i*_ and *a*_*ii*_ would act as an upper bound for growth parameters, and parameters could initially have values away from the trade-off curve (Shoval et al., 2012). Mechanistically, this upper bound could exist because of energetic and enzymatic constraints inside the microbe (Fuentes et al., 2021; Schink et al., 2022). This assumption reflects experimental conditions where a population may not be close to the trade-off curve in a new environment (Novak et al., 2006; Li et al., 2019). This view of trade-offs, where adaptation is limited externally by a trade-off function by mechanistic or energetic constraints, is in contrast to trade-offs being imposed through antagonistic pleiotropy between genes (Acerenza, 2016; Mauro and Ghalambor, 2020). The latter view typically focuses on the effects of a single genetic locus on different traits, often under different environmental conditions (Stearns, 2010). Since mutations in trait values are not constrained in early adaptation, populations may accumulate trait-increasing mutations in both *r*_*i*_ and *a*_*ii*_ faster than if (for example) mutations that increase *r*_*i*_ decreased *a*_*ii*_. Pleiotropic constraints such as these could help explain why some prior experimental studies have observed anti-correlations between growth and carrying capacity within experimental replicates, but nit across replicates (Novak et al., 2006). Pleiotropic constraints are therefore an excellent candidate for further study in the context of evolution on the invasion fitness landscape.

We also sought to study the effect of varying mutation supply parameters on the outcome of evolution. At lower mutation rates, lower proportions of trait-increasing mutations, and lower mutation effect sizes, evolution should proceed slowly and populations will take longer to reach their predicted ESS. Indeed, our calculation of the time taken to reach ESS under different starting conditions support this assumption (Appendix Section 5, Supplementary Figure 4). Thus, in studies that experimentally observe evolution of traits, whether populations evolve to their predicted states might depend on the timescales as well as the properties of the distribution of mutations. On the other hand, under mutation parameters relevant to microbial populations, evolution occurred fast enough in our simulations that populations were able to reach their predicted ESS under shorter time scales (Figure 3). Since estimates of mutation rates and distributions of mutations vary between different microbes (Denamur and Matic, 2006; Eyre-Walker and Keightley, 2007; Zhu et al., 2014; Ramiro et al., 2020), it would be important to take into account where specific populations lie on this spectrum for predictive purposes. However, further research into the number of sites genome-wide (and effect sizes of mutations) underlying growth and competition-related traits would be needed to fully parameterize such models.

In two-species simulations, we observe that certain combinations of mutation supply parameters lead to a destabilization of coexistence, contradicting the predictions from adaptive dynamics theory (Figures 4 and 6). This destabilization of coexistence depended on the initial values of growth parameters, as that influenced the evolutionary trajectory of populations, which sometimes led to a destabilization of coexistence even when the initial and final points of the trajectory were within the coexistence region (Figure 5). One hypothesis is that these two factors are connected through a progressive reduction in population size of one competitor, as populations unable to access enough large-effect, trait-increasing mutations become unable to maintain evolutionary trajectories that lead them back into the coexistence region. Such extinction spirals, where smaller populations get trapped in a positive feedback loop of decreasing population size, have been documented in many systems (Turkarslan et al., 2017; Zhao et al., 2018; Williams et al., 2021; Richards, 2024).

Finally, we only examined the mean values of *r*_*i*_ and *a*_*ii*_ in our evolving populations. This was mainly because our theoretical predictions indicated that populations would evolve towards a single trait value (the ESS). However, in simulations where multiple mutations competed with each other throughout the course of evolution, it is possible that there would be variation in the individual values of *r*_*i*_ and *a*_*ii*_, which could have implications for the trajectory of evolution, and in the two-species case, continued coexistence of both species. In experimentally evolved populations, population-level variance in growth rates has been observed (Novak et al., 2006; D’Souza and Kost, 2016; Gounand et al., 2016). It has also been well-established that the presence and maintenance of genetic variation influences the evolutionary trajectory of populations (Barton and Keightley, 2002; Barett and Schulter, 2008). Thus, differences in population level genetic variation that influence values of *r*_*i*_ and *a*_*ii*_ would have implications for how evolution would proceed.

In sum, it has long been appreciated that species interactions are important for determining cooccurrence patterns, and a large body of research shows that pairwise interactions can be sufficient to predict multi-species community assembly (Friedman et al., 2017; Venturelli et al., 2018; Dedrick et al., 2023; Lele et al., 2026). Invasion fitness landscapes have provided valuable insights into how these interactions may evolve, but mechanistic models of evolution can depart from these predictions in important ways, which may affect the interpretation of evolutionary experiments. Our results indicate several factors that influence the evolution and stable coexistence of species, such as the timescales of observation, the initial values of growth parameters, and the supply of mutations, in both intuitive and less obvious ways. Empirically parameterizing the supply of trait-altering mutations, their effect sizes, and pleiotropy between traits would allow for the integration of invasion fitness landscapes and population genetic models to better inform the interpretation of experimental data.

## Data availability statement

Data and code from this manuscript are available at https://github.com/kasturilele/adaptive-dynamics. This data and code will be placed in a public repository upon acceptance of this manuscript for publication.

