## Appendix 1 for "Fitness landscapes for species interactions: when do population genetics and adaptive dynamics diverge?"

#### 1 Summary

In sections 2, 3 and 4, we first tackle the single species case, where we obtain the fitness function from the growth equations, and then consider a tradeoff function between $r$ and $a$ where a candidate ESS is predicted. We then evaluate the properties of this ESS both graphically as well as analytically. Then, we extend this to a two-species case, where once again we assume that a similar tradeoff function exists between $r_i$ and $a_{ii}$ (that is, the single species parameters). We use this to explore parameter combinations where coexistence is predicted, and cases where evolution would destabilize coexistence. In section 5, we present the calculation for the time to fixation and the rate of fixation for the single-species case, using standard population genetics theory. In section 6, we present the supplementary figures and tables.

#### 2 Background

##### 2.1 Generalized Lotka-Volterra (gLV) model

The general form of the equation we are working with is:

$$\frac{dN_i}{dt} = (r_i + a_{ii}N_i + \sum_{j \neq i} a_{ij}N_j) * N_i \tag{1}$$

This equation has the growth rate on the left hand side, calculated in terms of the growth parameters. In this equation, $N_i$ represents the abundance of species $i$, $r_i$ is the basal growth rate, and $a_{ii}$ and $a_{ij}$ are coefficients that encompass all interactions each member of the focal species $i$ can have. $a_{ii}$ is the coefficient for intraspecific interaction (interaction of species $i$ with itself), and $a_{ij}$ are the coefficients for interspecific interaction (interaction of species $i$ with all other species $j$).

The conditions of coexistence under this framework are -

$$\rho < \frac{f_2}{f_1} < \frac{1}{\rho} \tag{2}$$

where $\rho$ and $\frac{f_2}{f_1}$ can be determined from the growth parameters as follows-

$$\begin{aligned} \rho &= \sqrt{\frac{a_{12} * a_{21}}{a_{11} * a_{22}}} \\ \frac{f_2}{f_1} &= \sqrt{\frac{a_{11} * a_{12}}{a_{21} * a_{22}}} * \frac{r_2}{r_1} \end{aligned} \tag{3}$$

(modified from Letten et al. (2017), also see the appendix of Lele et al. (2026)).

### 3 Mutant invading a population consisting of a single species

#### 3.1 Deriving the form of the fitness function

The growth equation for a single species ($N_r$) can be obtained from the general form of the equation as follows-

$$\frac{dN_r}{dt} = (r + aN_r) * N_r \tag{4}$$

We denote the parameters specific to the resident by marking them with $r$.

Now, a mutant ($N_m$) is introduced in this population. The growth equation for the mutant can be written as follows-

$$\frac{dN_m}{dt} = (r_m + a_m N_r) * N_m \tag{5}$$

We denote the mutant parameters as separate from the resident ones by marking them with $_m$. We make a few simplifying assumptions here. First, mutants are rare and low in frequency when they arise, so their growth depends on the presence of members of the resident species (hence the term $a_m N_r$) and the presence of mutants in the system will not initially affect the growth of the resident (hence the resident growth equations are unchanged).

This population will be at equilibrium when both resident and mutant growth rates are 0. This could happen either when $N_r = 0$ (trivial case that we do not need to consider further) or when $N_r = \frac{-r}{a}$ (note that this equilibrium will only be achieved for values of $a < 0$). Additionally, the mutant $N_m$ must always be at 0 at equilibrium.

Now, we identify the invasion fitness for the mutant and then figure out the parameter conditions for which a mutant would invade a stable population of the resident species. We can determine the direction of evolution by obtaining the stability matrix of the system of growth equations, and examining further the eigenvalue which contains the growth parameters for the mutant ($r_m$ and $a_m$).

The eigenvalue that contains the mutant parameters gives us the invasion fitness $s_m$.

$$s_m = r_m + a_m N_r \tag{6}$$

We note that $s_m$ is the per capita growth rate of the mutant. This aligns with our intuitive understanding that when the per capita growth rate is positive, i.e. when $s_m > 0$, the mutant will be able to invade the population. Substituting $N_r = \frac{-r}{a}$ in equation 6, from the equilibrium value we calculated from equation 4, we obtain-

$$s_m = r_m - \frac{r a_m}{a} \tag{7}$$

Thus, if $s_m$ is greater than 0, the mutant will be able to invade the existing population. Additionally, we can verify that when the mutant parameters are identical to the resident parameters, $s_m = 0$.

#### 3.2 Predicting the direction of evolution (finding and evaluating candidate ESS)

##### 3.2.1 $r_m$ and $a_m$ vary without constraint

We first consider the condition where there are no constraints over $r_m$ and $a_m$, and hence both these parameters are free to vary with mutations. However, since mutations are rare, and mutations in one parameter do not affect the other, we assume that only one of these will get mutated at a given time. Thus, we either have $r_m = r$ or $a_m = a$.

To understand in which direction $r$ or $a$ will evolve, we obtain the fitness gradient (the partial derivative with respect to the mutant parameters) at the resident trait value. For mutations of small effect, since mutant alleles will only be able to invade the population when $s_m > 0$, the sign of the derivative will tell us whether mutants with trait value greater or smaller than the resident trait value will be able to invade the population (see Otto and Day, 2007). So, when $a_m = a$, i.e. only $r$ varies:

$$\frac{\delta s_m}{\delta r_m} = 1 \tag{8}$$

Since equation 8 is always $> 0$, mutants with a growth rate $r_m$ can invade the population if $r_m > r$ and populations will continue to evolve higher values of $r$. Similarly, when $r_m = r$ i.e. only $a$ varies:

$$\frac{\delta s_m}{\delta a_m} = -\frac{r}{a} \tag{9}$$

Near the equilibrium, both $a_m$ and $a_r$ will be negative, and equation 9 is always $> 0$. Thus, mutants with interaction coefficient $a_m$ can invade the population if $|a_m| < |a|$, and the population will continue to evolve less negative values of $a$ (lower in magnitude).

The pairwise invasibility plots (PIPs) for these conditions give us the same information - that the population will eventually evolve towards $r_m > r$ and $|a_m| < |a|$. See supplementary figure 1.

This indicates that if there were no constraints on the evolution of $r$ and $a$, then the population will evolve towards higher values of $r$ and less negative values of $a$. This would lead to the value of $K$ becoming higher and higher, which is unsustainable. Thus, we henceforth assume the presence of some kind of trade-off function between $r$ and $a$.

#### 3.3 Adding constraints (tradeoff function) between the parameters

##### 3.3.1 First-order derivative equations

Consider the fitness function from the previous section (equation 7). Now, we consider what might happen if there was a function relating the parameters, $a = f(r)$. Specifically, we want to understand the fate of the mutation close to what would be a candidate ESS, i.e. a state where the resident population cannot be invaded by a mutant. The derivative of the fitness gradient with respect to the mutant gives us the direction in which evolution would proceed, so if we equate the derivative to 0, we obtain a condition under which mutants will not be able to invade.

$$s_m = r_m - \frac{r * f(r_m)}{f(r)} \tag{10}$$

Taking the derivative of $s_m$, we obtain-

$$\frac{\delta s_m}{\delta m}|_{(r_m=r_*)} = 1 - \frac{r_* * f'(r_*)}{f(r_*)} \tag{11}$$

Any function $f(r)$ that satisfies the condition $\frac{\delta s_m}{\delta m}|_{(r_m=r_*)} = 0$ for a value of $r_*$ will lead to a candidate ESS. Conversely, there are many possible tradeoff functions, for which no candidate ESS is predicted, and any value of $r_m$ either greater or lesser than the resident could invade the population.

##### 3.3.2 Second-order derivative equations

To determine whether this candidate ESS is evolutionary and convergence stable, we need to calculate the second order derivative with respect to the resident as well as the mutant. The criteria for different outcomes of evolutionary and convergence stability are given below (from Bowers and White, 2002; Diekmann, 2002)

| Properties of candidate ESS ( $\frac{\delta s_m}{\delta m}|_{(r_m=r_*)} = 0$ ) | conditions satisfied by second-order derivatives |
|---|---|
| Evolutionarily unbeatable strategy (EUS) | $D_m < 0$ |
| Convergence stable strategy (CS) | $D_r > D_m$ |
| Candidate ESS invades resident (EIR) | $D_r > 0$ |
| Mutually invadable (MI) | $D_r > -D_m$ |

Here, $D_r = \frac{\delta^2 s_m}{\delta r^2}|_{(r_m=r_*)}$ and $D_m = \frac{\delta^2 s_m}{\delta m^2}|_{(r_m=r_*)}$ .

We are mainly concerned with Conditions 1) and 2) here, as together these conditions can inform us whether the candidate ESS is, along with being an ESS, also convergence stable.

$$\frac{\delta^2 s_m}{\delta m^2}|_{(r_m=r_*)} = -\frac{r_* * f''(r_*)}{f(r_*)} \tag{12}$$

taking the second derivative of the fitness function with respect to the resident, we get-

$$\frac{\delta^2 s_m}{\delta r^2}|_{(r_m=r_*)} = \frac{2f'(r_*)}{f(r_*)} - \frac{2r_* f'(r_*)^2}{f(r_*)^2} + \frac{r_* * f''(r_*)}{f(r_*)} \tag{13}$$

since $\frac{\delta s_m}{\delta m}|_{(r_m=r_*)} = 0$ at $m = r = r_*$, we can simplify the second order derivative with respect to the resident as follows-

$$\frac{\delta^2 s_m}{\delta r^2}|_{(r_m=r_*)} = \frac{r_* * f''(r_*)}{f(r_*)} \tag{14}$$

Simplifying this in terms of the conditions mentioned above, a few conclusions immediately stand out-

1. No tradeoff function can satisfy the last condition to be mutually invadable, since $D_r = -D_m$.
2. Also, since $D_r = -D_m$, the second condition (convergence stability) will be satisfied for all $D_r > 0$, and never be satisfied if $D_r <= 0$.
3. Following from above, when $D_r > 0$ the candidate ESS will also be able to invade nearby resident strategies. This excludes us from having a case where an ESS is possible if we can reach it, but when a strategy is away from the ESS it cannot evolve towards it.
4. Since $D_r = -D_m$, the only two possible cases (where an ESS exists at all, and not just all r above or below resident being able to invade) are a stable equilibrium with EUS, CS and EIR, or an unstable equilibrium with none of these three possibilities.

In the next section, we will explore predictions for how r and alpha evolve assuming a certain shape of trade-off function.

#### 3.4 Predictions under specific constraint functions

When we substitute specific constraint functions with specific shapes in the invasion fitness equation, we can obtain predictions for convergence and evolutionary stability for them. Below, we explore whether the predicted ESS are convergence and evolutionarily stable for a specific function, and make predictions for how r and alpha will evolve under this function. Later in the paper, we will test these predictions with simulations, and see if they still hold if the assumptions of the adaptive dynamics framework are not fulfilled.

##### 3.4.1 Exponentially decreasing tradeoff between r and alpha

We consider a tradeoff between $r$ and $a$ such that $a$ decreases exponentially with $r$, such that $a = f(r) = C_1 * e^{C_2 * r}$. For different values of the constants $C_1$ and $C_2$, we can obtain obtain an ESS $r_*$ by setting $\frac{\delta s_m}{\delta m}|_{(r_m=r_*)} = 0$, and evaluate the properties of this ESS. Thus, we obtain $r_* = \frac{1}{C_2}$. We can also obtain the ESS graphically, from the PIP (figure 2 of text).

For our simulations, we consider the tradeoff function $a = f(r) = -0.0000206 * e^{3.2 * r}$, where we obtain a candidate ESS at $r = 0.3125$ from the PIP. We can verify this result by setting the derivative of the fitness gradient (i.e., equation 11) to 0, and obtaining the value of $r_*$ numerically, using Python.

Additionally, we can obtain the properties of this ESS by calculating the values of $D_r$ and $D_m$. For this tradeoff function, $D_r > 0$, $D_m < 0$, and thus this ESS is both convergence and evolutionarily stable. We can also verify this from the PIP. Thus, under this tradeoff function, populations will evolve towards the value of $r = 0.3125$.

### 4 Extending the single-species derivation to two species

#### 4.1 Deriving the form of the fitness function

The growth equation for two species ($N_{1r}$ and $N_{2r}$) can be obtained from the general form of the equation as follows-

$$\begin{aligned} \frac{dN_{1r}}{dt} &= (r_{1r} + a_{11r}N_{1r} + a_{12}N_{2r}) * N_{1r} \\ \frac{dN_{2r}}{dt} &= (r_{2r} + a_{22r}N_{2r} + a_{21}N_{1r}) * N_{2r} \end{aligned} \tag{15}$$

This equation is very similar to the earlier single-species equation, with the addition of the inter-specific interaction coefficients ($a_{12}$ and $a_{21}$) that represent the effect members of the other species have on the growth of the first species. As before, $_r$ indicates the resident trait value.

Now, a mutant ($N_{1m}$) is introduced in this population. We assume that only one species will have a mutant introduced in it at a given time. Since the equations are symmetric, we introduce a mutant in species 1 without loss of generality. The growth equation for this mutant can be written as follows-

$$\frac{dN_{1m}}{dt} = (r_{1m} + a_{11m}N_{1r} + a_{12}N_{2r}) * N_{1m} \tag{16}$$

We denote the mutant parameters as separate from the resident ones by marking resident parameters with $_r$ and mutant parameters with $_m$. Apart from the previous set of assumptions (i.e. mutations being rare and low frequency, such that the presence of mutants does not impact the growth of the resident), we additionally assume that mutations are not occurring in the inter-specific interaction coefficients.

This population (both species + mutant in species 1) will be at equilibrium when all growth rates are 0. Similar to the one-species case, the mutant $N_{1m}$ must always be at 0 at equilibrium. The equilibrium could be when both or one of $N_{1r}$ and $N_{2r}$ are 0, in which case, the model would simplify to the earlier one-species case. When both $N_{1r}$ and $N_{2r}$ are nonzero, we obtain the equilibrium values as follows-

$$\begin{aligned} N_{1r} &= \frac{r_{2r}a_{12} - r_{1r}a_{22r}}{a_{11r}a_{22r} - a_{12}a_{21}} \\ N_{2r} &= \frac{r_{1r}a_{21} - r_{2r}a_{11r}}{a_{11r}a_{22r} - a_{12}a_{21}} \end{aligned} \tag{17}$$

Similar to the previous section, we determine the invasion fitness $s_{1m}$ from the eigenvalue of the partial derivative of the system that contains the mutant parameters. $s_{1m}$ is once again the per capita growth rate of the mutant.

$$s_{1m} = r_{1m} + a_{11m}N_{1r} + a_{12}N_{2r} \tag{18}$$

Substituting the values of $N_{1r}$ and $N_2$ in the equation above, from the equilibrium value we calculated from equation 17, we obtain-

$$s_{1m} = r_{1m} + a_{11m}\frac{a_{12}r_{2r} - a_{22r}r_{1r}}{a_{22r}a_{11r} - a_{12}a_{21}} + \frac{a_{12}a_{21}r_{1r} - a_{11r}a_{12}r_{2r}}{a_{22}a_{11r} - a_{12}a_{21}} \tag{19}$$

Since the growth equations for species 1 and species 2 in this framework are symmetric, we can obtain a similar equation for the invasion fitness of a mutant in species 2.

$$s_{2m} = r_{2m} + a_{22m}\frac{a_{21}r_{1r} - a_{11r}r_{2r}}{a_{22r}a_{11r} - a_{12}a_{21}} + \frac{a_{12}a_{21}r_{2r} - a_{22r}a_{21}r_{1r}}{a_{22r}a_{11r} - a_{12}a_{21}} \tag{20}$$

Again, if $s_{1m}$ and $s_{2m}$ are greater than 0, the mutant will be able to invade the existing population. Additionally, we can verify that when the mutant parameters are identical to the resident parameters, $s_{1m} = 0$ and $s_{2m} = 0$.

#### 4.2 Predicting the direction of evolution (finding and evaluating candidate ESS)

Similar to the 1-species case, with no constraints on $r_1$, $a_{11}$, $r_2$, and $a_{22}$, they will evolve towards higher values of r and less negative values of alpha. We can verify this by obtaining the fitness gradient of the fitness function to understand the direction in which the parameters under selection will evolve. It is straightforward to observe that $\frac{\delta s_{1m}}{\delta r_{1m}} = 1$ and $\frac{\delta s_{2m}}{\delta r_{2m}} = 1$. By taking the partial derivative with respect to alpha, we obtain -

$$\begin{aligned} \frac{\delta s_{1m}}{\delta a_{11m}} &= \frac{a_{12}r_{2r} - a_{22r}r_{1r}}{a_{22r}a_{11r} - a_{12}a_{21}} \\ \frac{\delta s_{2m}}{\delta a_{22m}} &= \frac{a_{21}r_{1r} - a_{11r}r_{2r}}{a_{22r}a_{11r} - a_{12}a_{21}} \end{aligned} \tag{21}$$

However, to maintain coexistence, both resident and mutant parameters must satisfy the conditions given in equations 2 and 3. Simplifying those inequalities further, we get -

$$\begin{aligned} \frac{a_{21}}{a_{11}} &< \frac{r_2}{r_1} \\ \frac{a_{12}}{a_{22}} &< \frac{r_1}{r_2} \end{aligned} \tag{22}$$

From these inequalities, we further obtain -

$$
\begin{aligned}
a_{12}r_2 - a_{22}r_1 &> 0\\
a_{21}r_1 - a_{11}r_2 &> 0\\
a_{11}a_{22} - a_{21}a_{12} &> 0
\end{aligned} \tag{23}
$$

Thus, similar to the one-species case, we see that $\frac{\delta s_{1m}}{\delta a_{11m}} > 0$ and $\frac{\delta s_{2m}}{\delta a_{22m}} > 0$, given the constraint that $a_{11}$ and $a_{22}$ are always negative. Thus, even in this case, alphas will evolve towards less negative values in order to maintain coexistence. If coexistence is destabilized and one of the species goes extinct, then the remaining species' value of alpha will also evolve towards less negative values, according to the one-species case as detailed in the previous section.

From this, we conclude that mutations with no constraints in $r_1$, $a_{11}$, $r_2$, and $a_{22}$ will eventually lead to destabilization of coexistence, as $r_1$ and $r_2$ increase in magnitude and $a_{11}$ and $a_{22}$ decrease in magnitude (with no change in the value of $a_{12}$ and $a_{21}$). We can also verify this with our simulations, where one species eventually goes extinct as both populations accumulate mutations.

###### 4.2.1 First and second order partial derivatives under tradeoff functions

We now explore how the growth parameters evolve under different trade-off functions between these parameters. Substituting $a_{11} = f(r_1)$ and $a_{22} = g(r_2)$, we have-

$$
\begin{aligned}
s_{1m} &= r_{1m} + f(r_{1m})\frac{a_{12}r_{2r} - g(r_{2r})r_{1r}}{f(r_{1r})f(r_{2r}) - a_{12}a_{21}} + \frac{a_{12}a_{21}r_{1r} - f(r_{1r})a_{12}r_{2r}}{f(r_{1r})g(r_{2r}) - a_{12}a_{21}}\\
s_{2m} &= r_{2m} + f(r_{2m})\frac{a_{21}r_{1r} - f(r_{1r})r_{2r}}{f(r_{1r})g(r_{2r}) - a_{12}a_{21}} + \frac{a_{12}a_{21}r_{2r} - g(r_{2r})a_{21}r_{1r}}{f(r_{1r})g(r_{2r}) - a_{12}a_{21}}
\end{aligned} \tag{24}
$$

like in the previous section, we calculate the derivative of the fitness function 24 to find the candidate ESS-

$$
\begin{aligned}
\frac{\delta s_{1m}}{\delta r_{1m}}|_{(r_{1m}=r_{1*})} &= 1 + \frac{(r_{2*}a_{12} - f(r_{2*})r_{1*}) * f'(r_{1*})}{f(r_{1*}f(r_{2*}) - a_{12}a_{21}}\\
\frac{\delta s_{2m}}{\delta r_{2m}}|_{(r_{2m}=r_{2*})} &= 1 + \frac{(r_{1*}a_{21} - f(r_{1*})r_{2*}) * f'(r_{2*})}{f(r_{1*}f(r_{2*}) - a_{12}a_{21}}
\end{aligned} \tag{25}
$$

Like the single-species case discussed above, any function that satisfies $\frac{\delta s_m}{\delta m}|_{(r_m=r_*)} = 0$ will lead to a candidate ESS, and we can evaluate the properties of this candidate ESS by calculating the value of the second order partial derivatives with respect to the resident and the mutant.

In order to ensure that both species coexist at the candidate ESS, we also impose the additional condition that both species' equilibrium population sizes (given in equation 17) be greater than 0. If a candidate ESS does not satisfy this condition, then coexistence will get destabilized as both species evolve in the direction of the ESS. We can numerically solve the combined partial differential equations for specific tradeoff functions to obtain conditions under which the all the above conditions are true, and conditions under which evolving towards the candidate ESS would lead to destabilization of coexistence.

Now, we must take the second order partial derivatives for both fitness functions, with respect to the resident and mutant parameters. The second order derivative with respect to the mutant parameters is straightforwardly obtained from the first order partial derivative-

$$
\begin{aligned}
D_{m1} &= \frac{(r_{2*}a_{12} - f(r_{2*})r_{1*}) * f''(r_{1*})}{f(r_{1*}f(r_{2*}) - a_{12}a_{21}}\\
D_{m2} &= \frac{(r_{1*}a_{21} - f(r_{1*})r_{2*}) * f''(r_{2*})}{f(r_{1*}f(r_{2*}) - a_{12}a_{21}}
\end{aligned} \tag{26}
$$

Taking the second derivative with respect to the resident trait value, we get the following equation for species 1-

$$\frac{\delta^2 s_{1m}}{\delta r_1^2}|_{(r_{1m}=r_{1*})} = \frac{2g(r_2)^2 f(r_{1*})f'(r_{1*})}{g(r_2)f(r_{1*}) - a_{12}a_{21})^2} + \frac{2g(r_2)^2(a_{12}a_{21}r_{1*} - r_{2*}a_{12}f(r_{1*}))f'(r_{1*})^2}{(g(r_2)f(r_{1*}) - a_{12}a_{21})^3}$$
$$+ \frac{2g(r_2)^2(r_{2*}a_{12} - g(r_2))f(r_{1*})f'(r_{1*})^2}{(g(r_2)f(r_{1*}) - a_{12}a_{21})^3} - \frac{2g(r_2)f'(r_{1*})(a_{12}a_{21} - r_{2*}a_{12}f'(r_{1*}))}{(g(r_2)f(r_{1*}) - a_{12}a_{21})^3} - \frac{r_{2*}a_{12}f''(r_{1*})}{g(r_2)f(r_{1*}) - a_{12}a_{21}}$$
$$- \frac{g(r_2)(a_{12}a_{21}r_{1*} - r_{2*}a_{12}f(r_{1*}))f''(r_{1*})}{(g(r_2)f(r_{1*}) - a_{12}a_{21})^2} - \frac{g(r_2)(r_{2*}a_{12} - g(r_2)r_{1*})f''(r_{1*})}{(g(r_2)f(r_{1*}) - a_{12}a_{21})^2} \quad (27)$$

We can simplify this equation by grouping all the terms that contain $f''(r_{1*})$, and all the terms that contain $f'(r_{1*})$. Further, we use equation 25 to simplify the grouped terms that contain $f'(r_{1*})$, and observe that all terms except the ones that contain $f''(r_{1*})$ add up to 0. We obtain the following expression after simplification-

$$D_{r1} = -\frac{(r_{2*}a_{12} - f(r_{2*})r_{1*}) * f''(r_{1*})}{f(r_{1*})f(r_{2*}) - a_{12}a_{21}} \quad (28)$$

Since the equations for species 2 are similar to the equations for species 1, we obtain a similar expression for $D_{r2}$ -

$$D_{r2} = -\frac{(r_{1*}a_{21} - f(r_{1*})r_{2*}) * f''(r_{2*})}{f(r_{1*})f(r_{2*}) - a_{12}a_{21}} \quad (29)$$

Like the one-species case, we once again observe that $D_{r1} = -D_{m1}$ and $D_{r2} = -D_{m2}$, and we can draw similar conclusions about the properties of the candidate ESS as we did in the section above.

##### 4.3 Finding the ESS and evaluating predictions of coexistence under specific tradeoff functions

Similar to the one-species case, we consider an exponentially decreasing tradeoff function for both species. Thus, we have $a_{11} = f(r_1) = C_1 * e^{C_2 * r_1}$ and $a_{22} = g(r_2) = C_3 * e^{C_4 * r_2}$. For different values of the constants $C_1$, $C_2$, $C_3$, and $C_4$, we can once again obtain the values of $r_{1*}$ and $r_{2*}$ (the candidate ESS), and check whether this candidate ESS is convergence and evolutionarily stable both graphically as well as numerically.

Having shown that a given pair of tradeoff functions leads to a candidate ESS that is both convergence and evolutionarily stable, we check whether populations coexist at the candidate ESS. We use the framework outlined by Chesson and others that we adapted for our equations in our previous work to obtain predictions of coexistence at the ESS. Based on the initial parameter values and the values of the constants used in the tradeoff functions, we have two possible outcomes-

1. Both populations coexist even after reaching their respective ESS.
2. One population goes extinct and the other population reaches its single-species ESS.

For our simulations, we consider the set of tradeoff functions $a_{11} = f(r_1) = -0.0000206 * e^{3.2 * r_1}$, $a_{22} = f(r_2) = -0.0000108 * e^{3.5 * r_2}$. We obtain a candidate ESS at $r_1 = 0.498344$ and $r_2 = 0.362690$ numerically by setting the derivative of the fitness gradient (i.e., equation 25) to 0, and obtaining the value of $r_{1*}$ and $r_{2*}$. We verify this ESS graphically using PIPs (main text figure). Additionally, we can obtain the properties of the candidate ESS by calculating the values of $D_{r1}$, $D_{r2}$, $D_{m1}$, and $D_{m2}$. For these tradeoff functions, $D_{r1}, D_{r2} > 0$, $D_{m1}, D_{m2} < 0$, and thus this ESS is both convergence and evolutionarily stable. We can also verify this from the PIPs. Thus, under these tradeoff functions, populations will evolve towards the value of $r_1 = 0.498344$ and $r_2 = 0.362690$.

For these specific tradeoff functions, we numerically obtain predictions of coexistence when both species reach their predicted ESS using the formula outlined above (equation 3). For the functions $a_{11} = f(r_1) = -0.0000206 * e^{3.2 * r_1}$, $a_{22} = f(r_2) = -0.0000108 * e^{3.5 * r_2}$, species are predicted to coexist at the ESS. We obtain additional predictions of coexistence for a range of values for $C_1$, $C_2$, $C_3$, and $C_4$ in the main text.

#### 5 Calculating the transit time along a tradeoff curve

In this section, we calculate the (state-dependent) rate of fixation for mutations in $r$. Following standard population genetics theory, we suppose that the probability of fixation is given by $2s$, where $s$ is the invasion fitness (Eqn. 3). When evolution is constrained by a tradeoff function of the form $a_{11} = C_1 e^{C_2 r_1}$, we can replace $r_{1m}$ and $a_{11m}$ in eqn. 7 with these values, to obtain

$$s = (r + \Delta r) - \frac{rC_1 e^{C_2(r+\Delta r)}}{rC_1 e^{C_2 r}} = \Delta r + r(1 - e^{C2\Delta r}). \tag{30}$$

Here, $\Delta r$ is the effect size of a mutation that alters the growth rate, $r$. Supposing a genome-average mutation rate $\mu_{r_+}$ for $r$-increasing mutations and genome length $L$, we can then calculate the expected rate of fixation of $r$-increasing mutations ($f_{r_+}$) per generation as

$$\mathbb{E}[f_{r_+}] = 2LN_{\text{eq}}\mu_{r_+}s = 2L\mu_{r_+}\left(re^{-C_2 r}\right)\left(\frac{\Delta r + r(1 - e^{C2\Delta r})}{C_1}\right). \tag{31}$$

This equation incorporates the expected values of population size $N_{\text{eq}}$ and $s$, both of which are dependent on the distance from the ESS and therefore time-dependent in the model. The expected rate of change in $r$ per generation can be found by taking the product $\mathbb{E}[f_{r_+}]\Delta r$. To approximate the expected time to reach some fine value of $r_f$ from a given initial value of $r_i$, we take the integral

$$\mathbb{E}[t_{r_f \leftarrow r_i}] \approx \int_{r_i}^{r_f} \frac{1}{\mathbb{E}[f_{r_+}]\Delta r} dr, \tag{32}$$

where $r_i$ is the value of $r$ at the ESS. The value of $\mathbb{E}[t_{r_f}]$ is only approximate, because we use the expected value of the rate of fixation in the denominator rather than considering the full distribution of possible times to fixation, and because we do not include the time required for a mutation to transit from low frequency to fixation. This integral can be evaluated in `Mathematica` to obtain

$$\mathbb{E}[t_{r_f} \leftarrow t_{r_i}] = \frac{-C_1}{2L\mu_{r_+}(\Delta r)^2}\left(W(s_{r_f}, C_2, \Delta r) - W(s_{r_i}, C_2, \Delta r)\right), \tag{33}$$

where $W(r, C_2, \Delta r)$ is defined as

$$W(r, C_2, \Delta r) = \text{Ei}[C_2 r] - e^{\frac{C_2\Delta r}{-1+e^{C_2\Delta r}}}\,\text{Ei}\left[\frac{-C_2 s_r}{-1 + e^{C_2\Delta r}}\right], \tag{34}$$

where $s_r$ is the invasion fitness at a given value of $r$, $s_r = \Delta r + r - re^{C_2\Delta r}$, and Ei$[z]$ is exponential integral, $\text{Ei}[z] = -\int_{-z}^{\infty} e^{-t}/t \; dt$.

We validated our calculation by comparing expected values of $\mathbb{E}[t_{r_f} \leftarrow t_{r_i}]$ to values obtained from stochastic simulation, as well as $r$-dependent fixation rates. Our expression for $r$-dependent fixation rates is obtained by taking the product of the genome-wide rate of mutation ($p_B L\mu$), the equilibrium population size ($-{}^{r}/_{a}$), and twice the invasion fitness at the given value of $r$. In these simulations, $\Delta r$ is fixed to a given value (ranging between 0.02 and 0.07), such that each effect mutation is of the same magnitude. We find close agreement between our predictions and quantities obtained by averaging over $10^3$ stochastic simulations (Fig. S4).

We also investigated how $\left(W(s_{r_f}, C_2, \Delta r) - W(s_{r_i}, C_2, \Delta r)\right)$ scales as function of $\Delta r$. If $\left(W(s_{r_f}, C_2, \Delta r) - W(s_{r_i}, C_2, \Delta r)\right)$ is relatively constant as a function of $\Delta r$, then the impact of effect sizes $\Delta r$ on the time to reach the ESS will be dominated by the term in the denominator of eqn. . We found that $\left(W(s_{r_f}, C_2, \Delta r) - W(s_{r_i}, C_2, \Delta r)\right)$ varies only modestly as a function of $\Delta r$. When $r_f$ is close to the ESS (*i.e.*, larger values of $r_f$ in Fig. S4), there is a peak in $\left(W(s_{r_f}, C_2, \Delta r) - W(s_{r_i}, C_2, \Delta r)\right)$ near $2({}^{1}/_{C_2} - r_f)$. This is expected because the ESS occurs at $r = {}^{1}/_{C_2}$ in the single-species model – the invasion fitness for mutations with large effects decreases as we approach the ESS (in part they can overshoot the ESS if they are large enough in magnitude, resulting in diminishing fitness improvements). Even with this increase in the time to reach the ESS for large-effect alleles, the scaling of ${}^{1}/_{\Delta r^2}\left(W(s_{r_f}, C_2, \Delta r) - W(s_{r_i}, C_2, \Delta r)\right)$ as a function of $\Delta r$ is dominated by the term ${}^{1}/_{\Delta r^2}$ (Fig. S4E), suggesting that the time to reach the ESS is indeed dominated by the ${}^{1}/_{\Delta r^2}$ term in the denominator.

### 6 Supplemental Figures and Tables

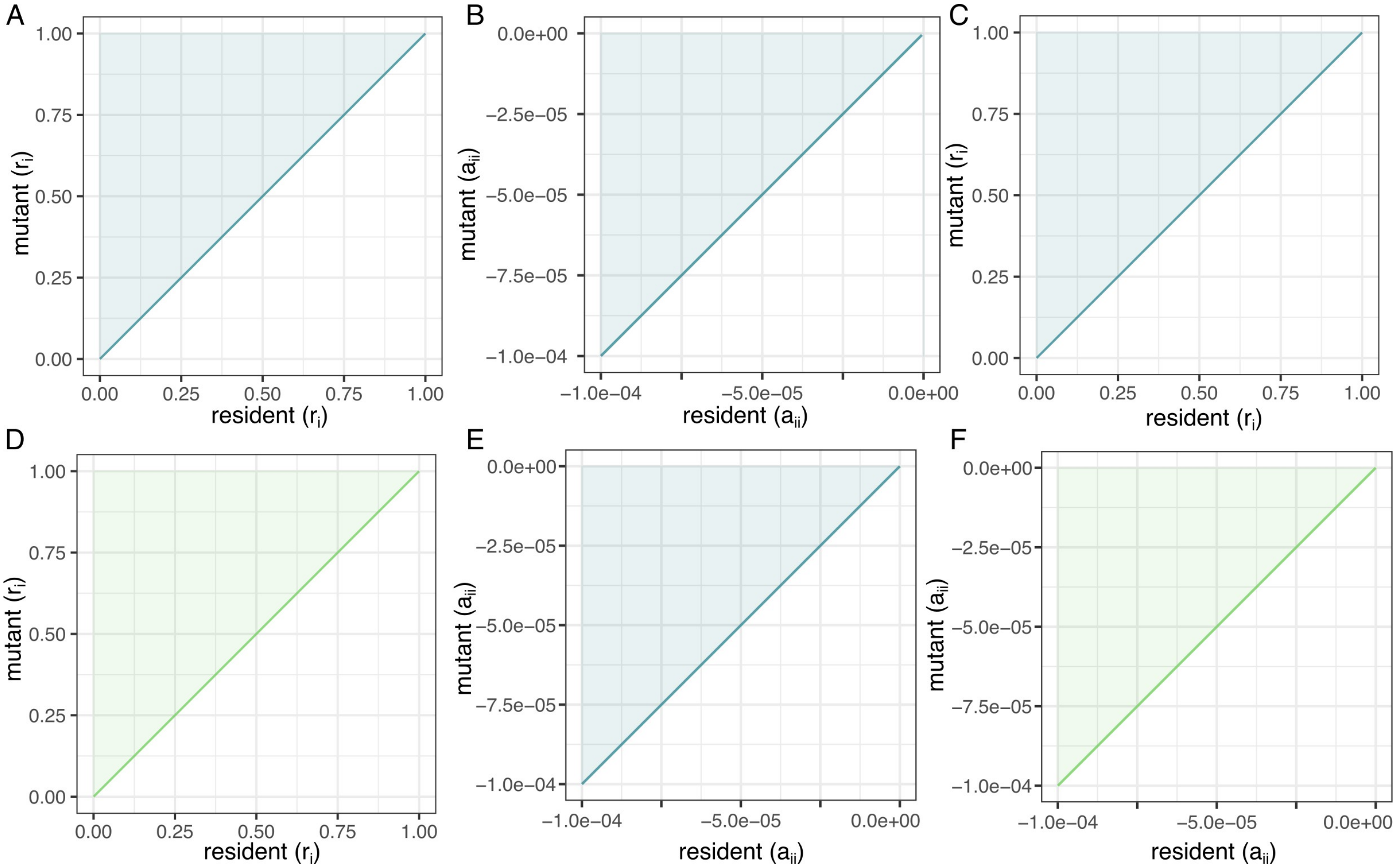


**Supplementary figure 1. Theoretical predictions of evolution without tradeoffs, where mutations accumulate separately in $r_i$ and $a_{ii}$.** A. Pairwise invasibility plot for $r_1$ for single species, showing that mutants can invade when they are bigger than the resident. B. Pairwise invasibility plot for $a_{11}$ for single species, showing that mutants can invade when less negative than the resident. C. Pairwise invasibility plot for $r_1$ for two species. D. Pairwise invasibility plot for $r_2$ for two species. C and D show that mutants can invade when they are bigger than the resident. E. Pairwise invasibility plot for $a_{11}$ for two species. F. Pairwise invasibility plot for $a_{22}$ for two species. E and F show that mutants can invade when less negative than the resident.

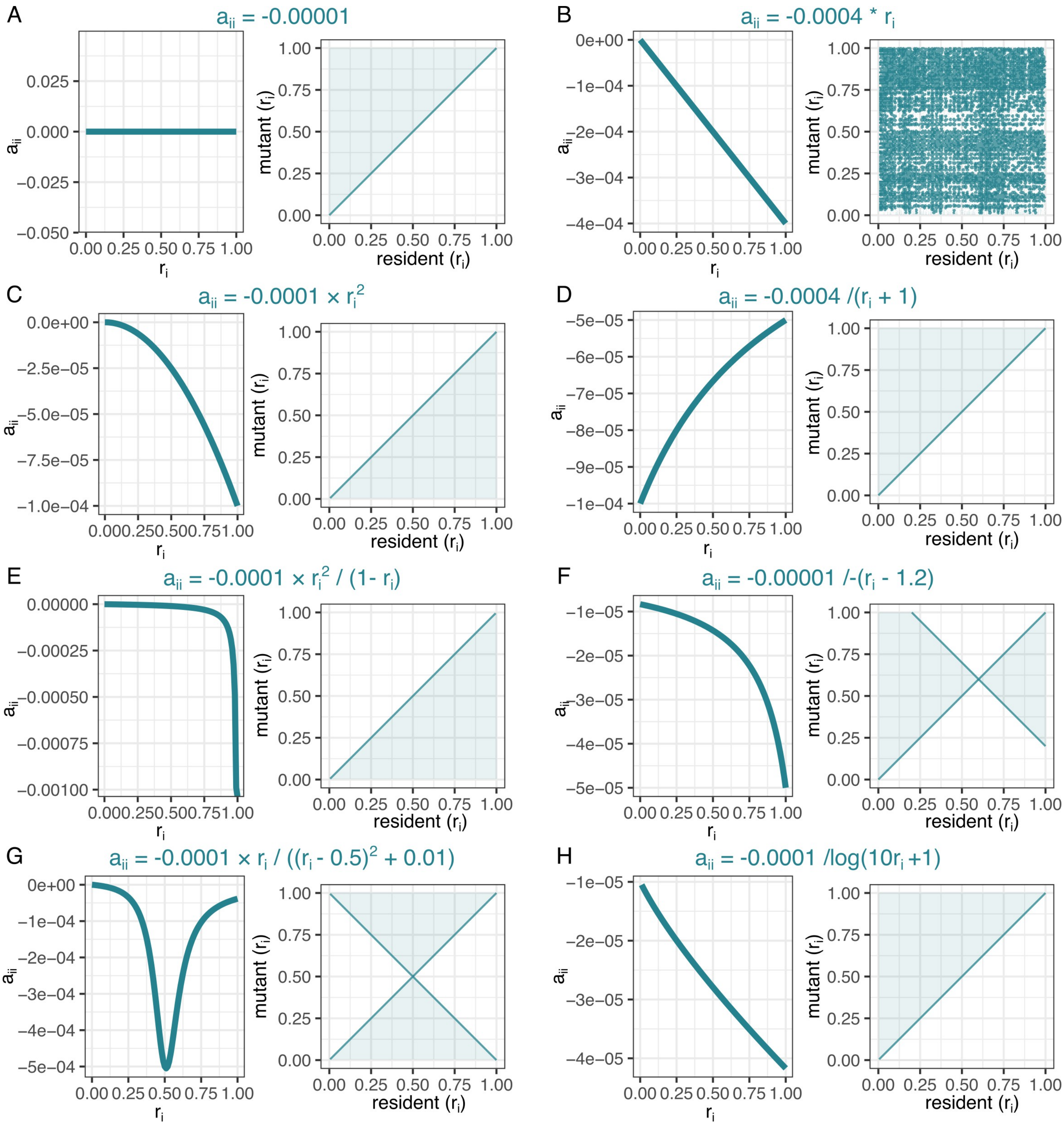


**Supplementary figure 2. Theoretical predictions of evolution for various tradeoff functions between $r_i$ and $a_{ii}$.** In each panel, the tradeoff function is on the left side and the pairwise invasibility plot is on the right side. A. $a_{ii} = C_1$, no tradeoff between $r_i$ and $a_{ii}$. B. $a_{ii} = C_1 * r_i$, linear tradeoff. C. $a_{ii} = C_1 * r_i^2$. D. $a_{ii} = \dfrac{C_1}{r_i + C_2}$. E. $a_{ii} = \dfrac{C_1 * r_i^2}{C_2 - r_i}$. F. $a_{ii} = \dfrac{C_1}{r_i - C_2}$. G. $a_{ii} = \dfrac{C_1 * r_i}{(r_i - C_2)^2 + C_3}$. H. $a_{ii} = \dfrac{C_1}{\log(C_2 * r_i + C_3)}$.

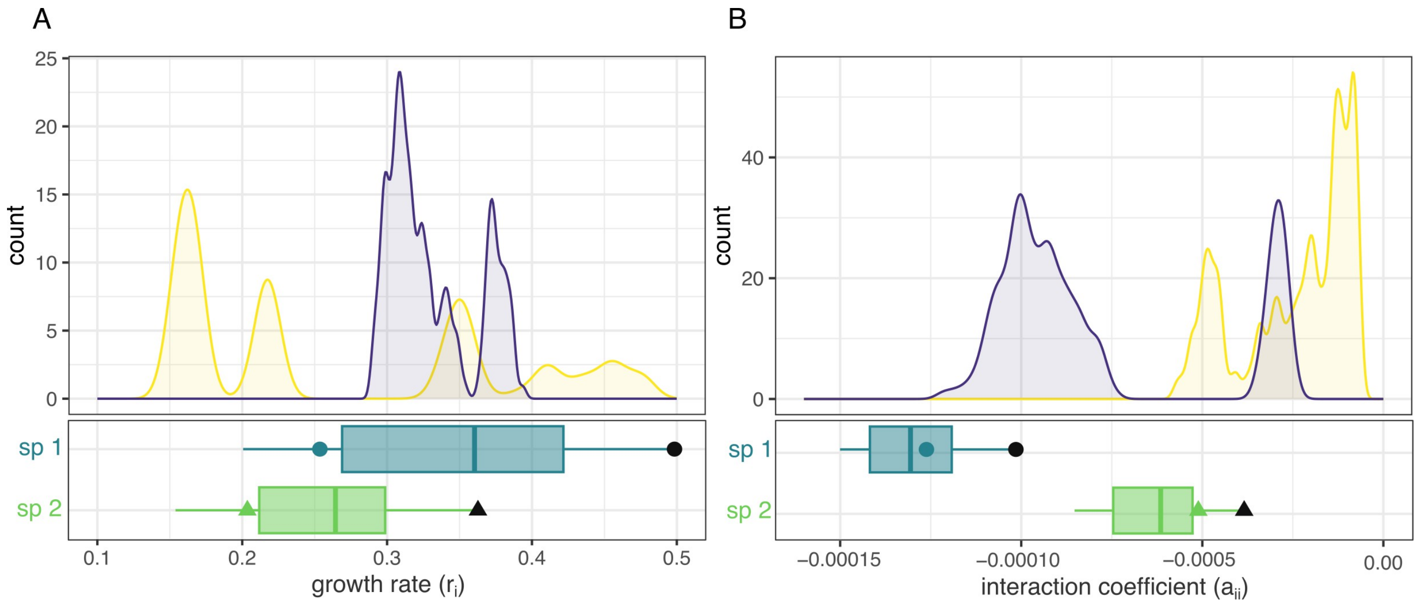


**Supplementary figure 3. Distributions of initial values of $r_i$ and $a_{ii}$ for all simulations.** A. Density plots showing the distribution of $r_i$ values, experimentally measured from sourdough microbes in our previous study (Lele et al., 2026). B. Density plots showing the distribution of $a_{ii}$ values from the same study, scaled by a factor of $10^3$ . In both plots, the yellow distribution indicates parameters measured from bacteria and dark blue distribution indicates parameters measured from yeast. The box plots below indicate the distribution of parameters for simulations where multiple sets of initial parameters were used (main text Figure 5). Colored circle and triangle indicate parameters used in all other simulations (where only a single set of initial parameters was used). The black circle and triangle indicates parameter values at ESS predicted under tradeoff functions (main text Figure 2).

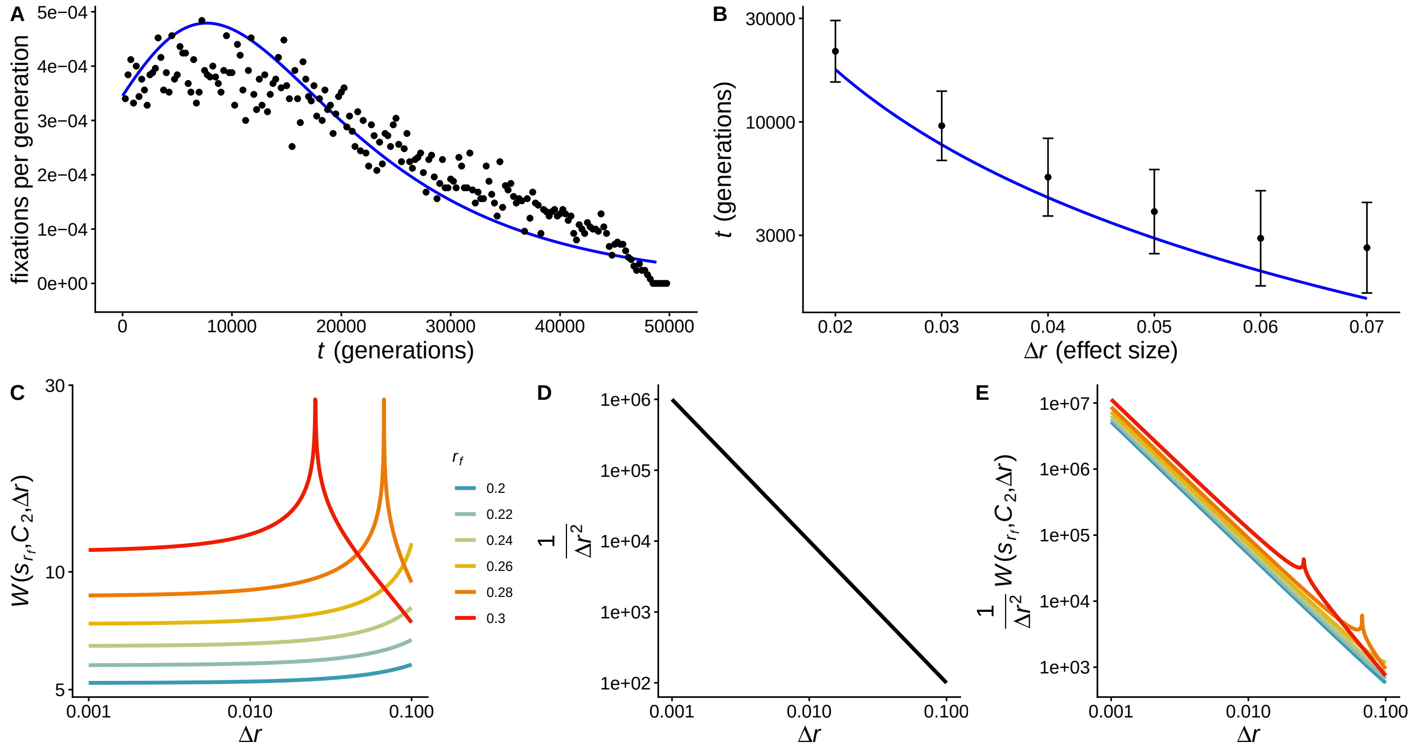


**Supplementary figure 4. Time to transit from $r_i$ to $r_f$ along a tradeoff curve** A. Expected rate of substitution per generation (blue line) as compared to the observed mean over $10^3$ simulations (black dots) B. Time to transit from $r_i = 0.05$ to $r_f = 0.2$ for a range of effect sizes. Points show the mean and standard deviation from 1,000 simulations, while blue line shows the expectation. C. $W(s_{r_f}, C_2, \Delta r) - W(s_{0.01}, C_2, \Delta r)$ as a function of $\Delta r$ for a range of values of $r_f$. D. $\frac{1}{\Delta r^2}$ as a function of $\Delta r$. E. $\frac{1}{\Delta r^2}\left(W(s_{r_f}, C_2, \Delta r) - W(s_{0.01}, C_2, \Delta r)\right)$ as a function of $\Delta r$ for the same values of $r_f$ as in C. *Additional parameters in A,B:* $L = 10^3$, $\mu = 2e-8$, $C_1 = -2.06e-5$, $p_B = 0.25$.

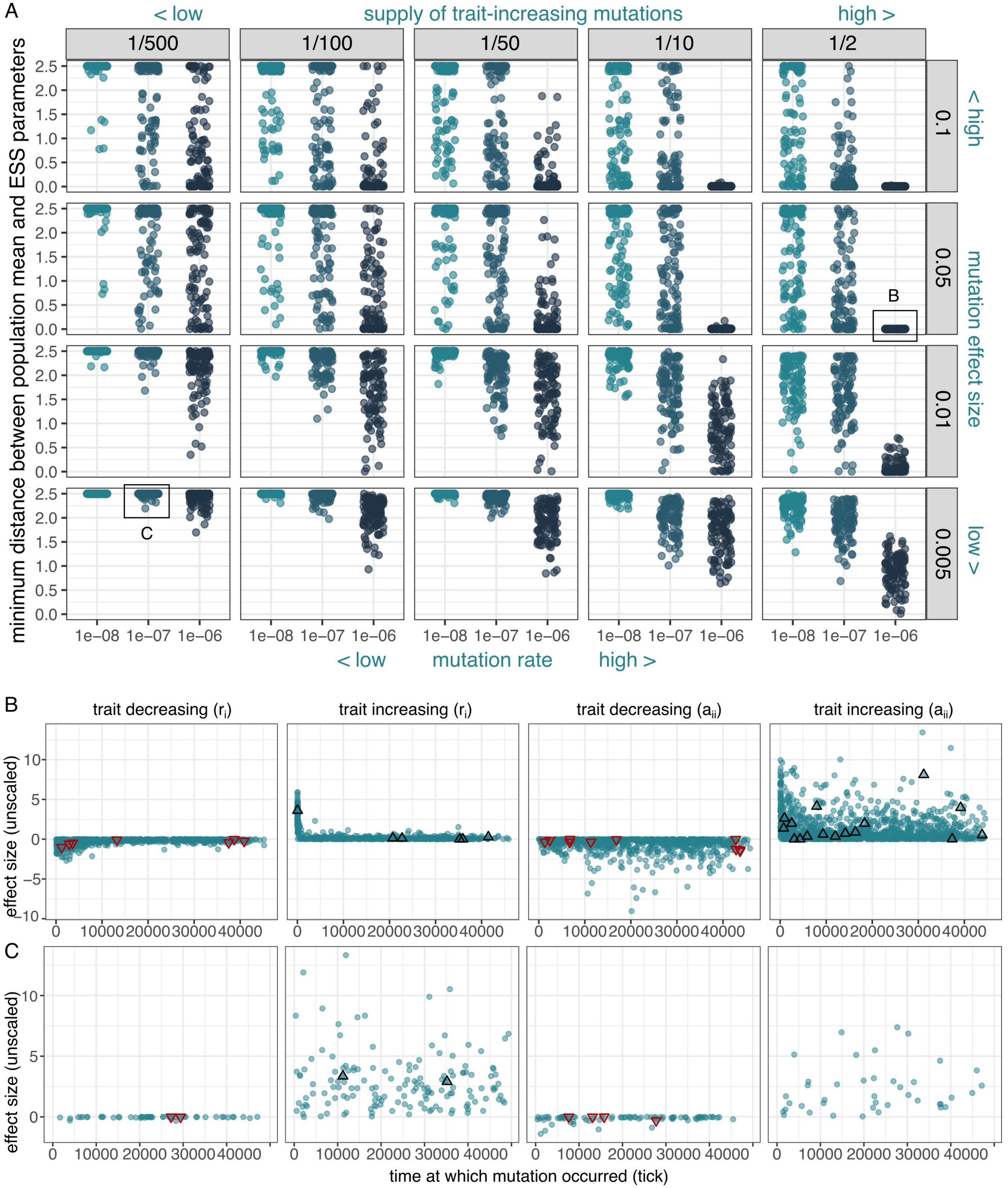


**Supplementary figure 5. Additional simulation results for single-species evolution over 50,000 ticks.** A. Minimum distance between the population mean growth parameters and the ESS parameters for different mutation rates ($\mu$), proportion of beneficial mutations ($m_k$), and mutation effect sizes ($mutr$ and $muta$). Colors indicate different mutation rates (also on x axis). B. Time of origin vs effect size (not scaled by the multiplier $\beta_r$ and $\beta_a$) for all fixed mutations from simulations with $mut = 10^{-6}$, $m_k = 1/2$, $mutr = 0.05$ and $muta = 0.000005$. C. Time of origin vs effect size for all fixed mutations from simulations with $\mu = 10^{-7}$, $m_k = 1/500$, $\beta_r = 0.005$ and $\beta_a = 0.0000005$. Dots indicate all fixed mutations, and red and black triangles indicate mutations fixed in the trajectory highlighted in the main figure.

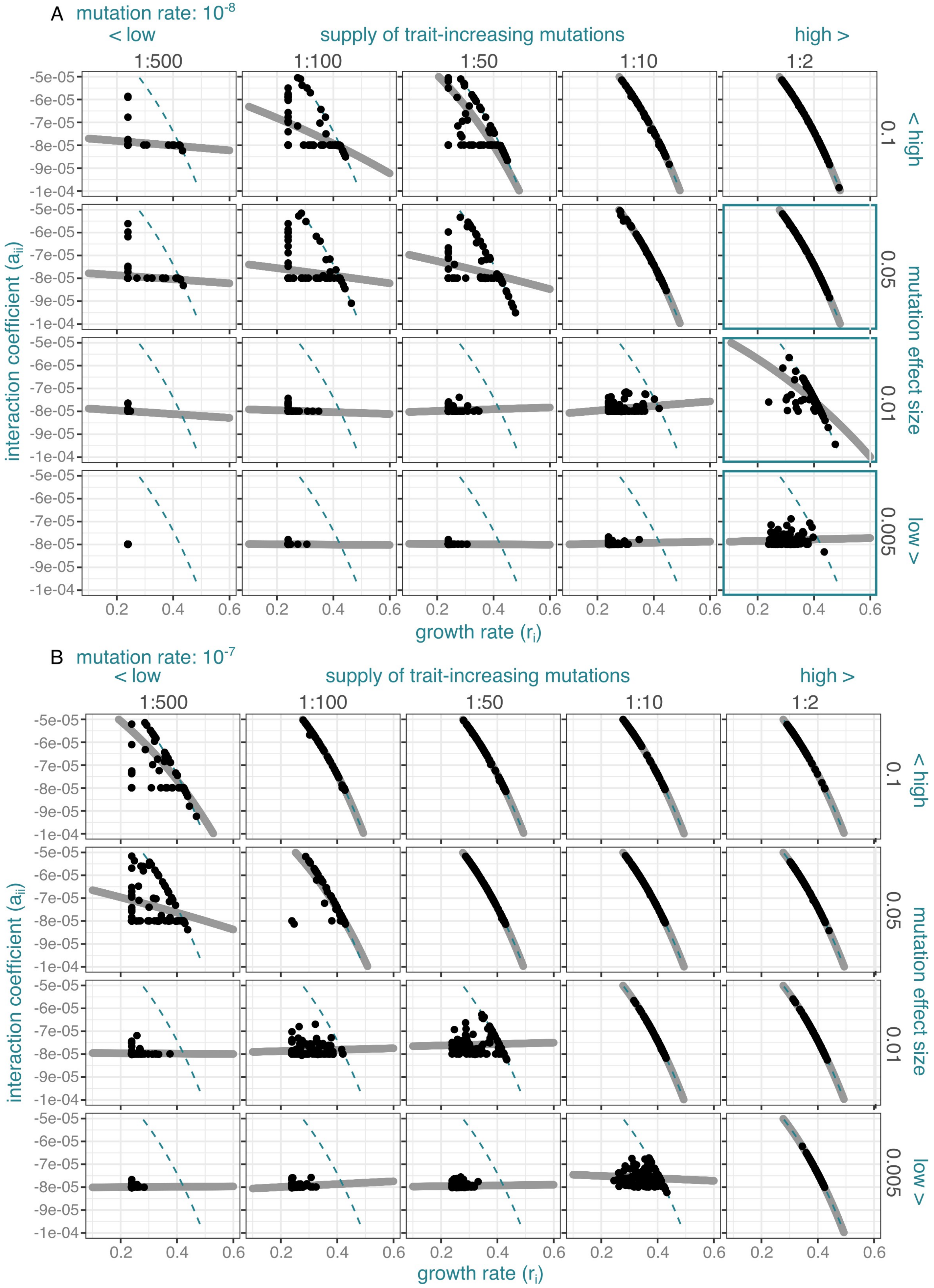
A mutation rate: $10^{-8}$
< low
supply of trait-increasing mutations
high >
1:500
1:100
1:50
1:10
1:2
< high
mutation effect size
low >
0.1
0.05
0.01
0.005
interaction coefficient ($a_{ii}$)
-5e-05
-6e-05
-7e-05
-8e-05
-9e-05
-1e-04
0.2
0.4
0.6
growth rate ($r_i$)
B mutation rate: $10^{-7}$
< low
supply of trait-increasing mutations
high >
1:500
1:100
1:50
1:10
1:2
< high
mutation effect size
low >
0.1
0.05
0.01
0.005
interaction coefficient ($a_{ii}$)
growth rate ($r_i$)

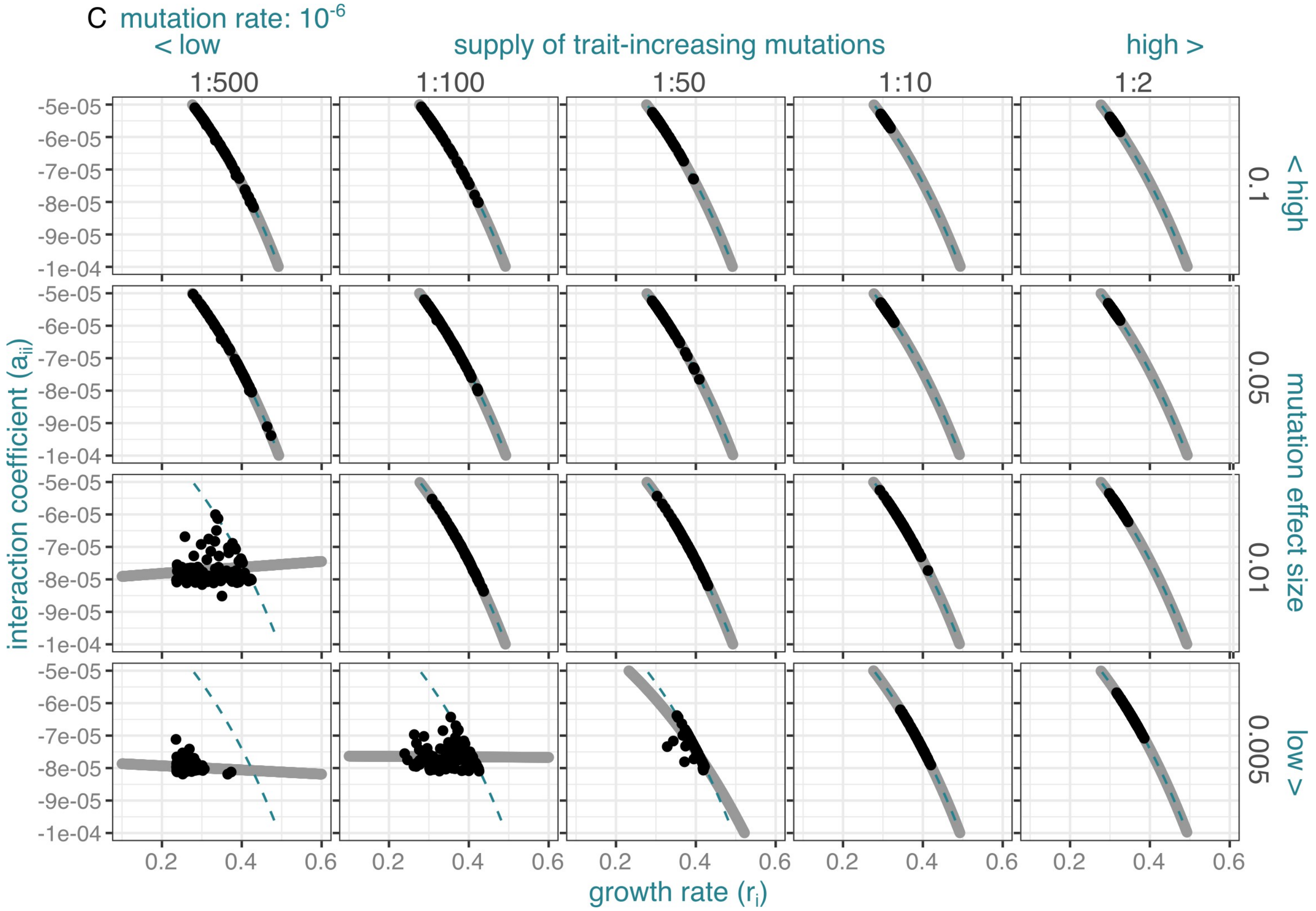


**Supplementary figure 6. Tradeoff functions fitted to simulated data for all single-species simulations.** Black dots indicate population mean values of $r_1$ and $a_{11}$ at 50,000 ticks, blue dashed line indicates original tradeoff function used in simulations, and grey line indicates tradeoff function fitted to the data. Blue square outlines indicate results shown in main text figure 3 D,E,F.

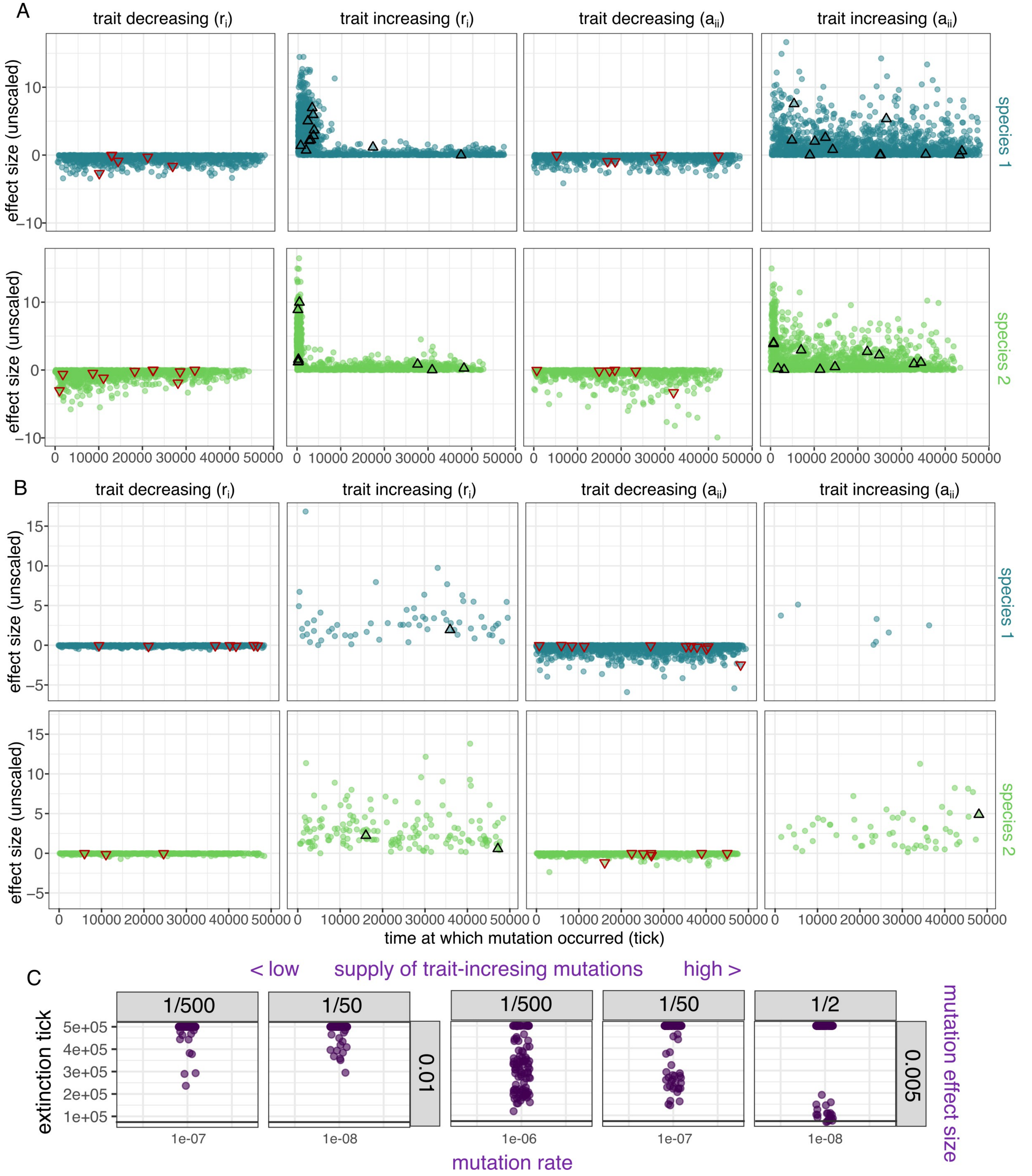


**Supplementary figure 7. Additional simulation results for two-species evolution over 50,000 ticks.** A. Time of origin vs effect size (not scaled by the multiplier $\beta_r$ and $\beta_a$) for all fixed mutations from simulations with $\mu = 10^{-6}$, $m_k = 1/2$, $\beta_r = 0.01$ and $\beta_a = 0.000001$. B. Time of origin vs effect size for all fixed mutations from simulations with $\mu = 10^{-6}$, $m_k = 1/500$, $\beta_r = 0.005$ and $\beta_a = 0.0000005$. Colors indicate species in which mutations arose, dots indicate all fixed mutations, and red and black triangles indicate mutations fixed in the trajectory highlighted in the main figure. C. Time at which one population went extinct in individual simulation runs (n=100) when simulations ran over longer time scales (500,000 ticks) for a few representative values of $\mu$, $m_k$, $\beta_r$ and $\beta_a$, showing that even for lower supplies of mutations, some populations eventually went extinct. Dark gray line indicates the time at which the simulations in the main text terminated (50,000 ticks).

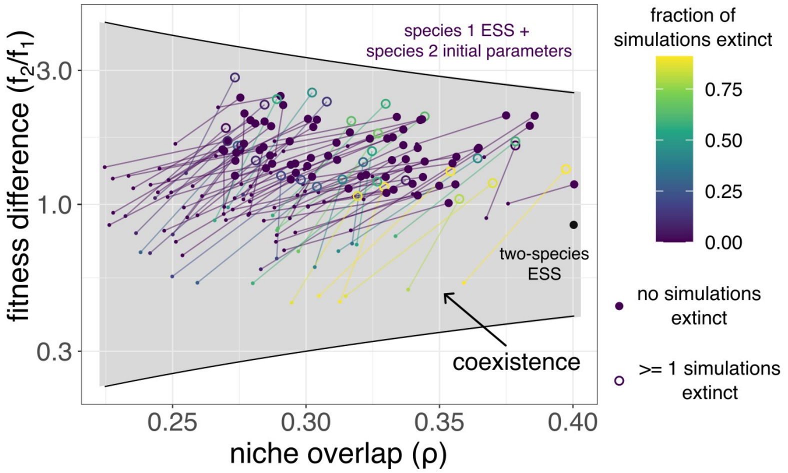


**Supplementary figure 8. Additional predictions of coexistence and proportions of replicates extinct in simulations.** A. Same as main text figure 2, shown here for comparison. B. Predictions of coexistence between 100 randomly chosen initial parameter combinations of two species, along with predictions for species 1 ESS and species 2 initial parameters. Open circles indicate parameter combinations which had >= 1 run of simulation go extinct. Colors indicate fraction of simulations that went extinct after 50,000 generations. Figure shown here to illustrate that for this combination, predictions of coexistence do not lie close to or outside the boundary of the region of coexistence.

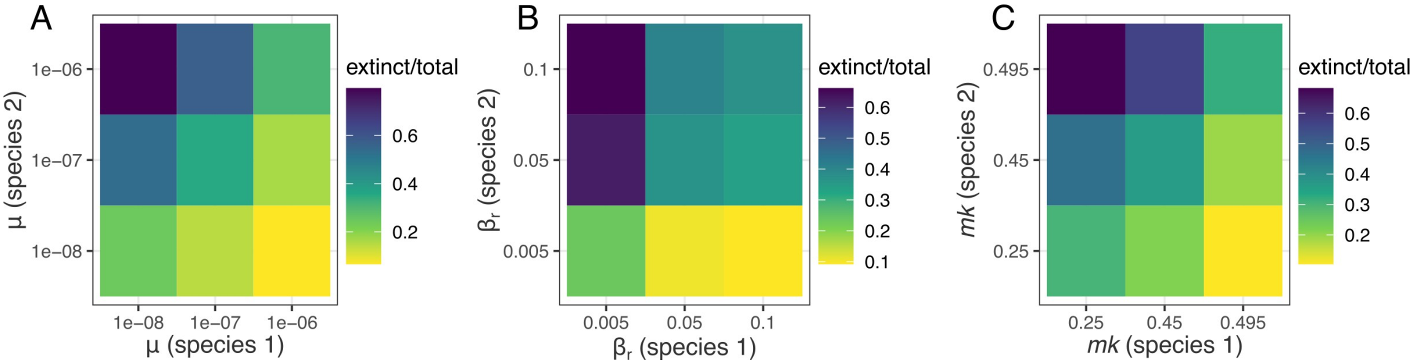


**Supplementary figure 9. Preliminary analyses of extinctions in simulations where the supply of mutations varied between populations.** A. Proportions of simulations in which one species went extinct for simulations grouped by mutation rate. B. Proportions of simulations in which one species went extinct for simulations grouped by mutation effect size multiplier. C. Proportions of simulations in which one species went extinct for simulations grouped by supply of trait-increasing mutations.

| ID | $m_k$ | $\beta_r$ | $\beta_a$ | $\mu$ | | ID | $m_k$ | $\beta_r$ | $\beta_a$ | $\mu$ |
|---|---|---|---|---|---|---|---|---|---|---|
| 1 | 1/2 | 0.005 | 5.00E-07 | 1.00E-08 | | 31 | 1/2 | 0.05 | 5.00E-06 | 1.00E-08 |
| 2 | 1/10 | 0.005 | 5.00E-07 | 1.00E-08 | | 32 | 1/10 | 0.05 | 5.00E-06 | 1.00E-08 |
| 3 | 1/50 | 0.005 | 5.00E-07 | 1.00E-08 | | 33 | 1/50 | 0.05 | 5.00E-06 | 1.00E-08 |
| 4 | 1/100 | 0.005 | 5.00E-07 | 1.00E-08 | | 34 | 1/100 | 0.05 | 5.00E-06 | 1.00E-08 |
| 5 | 1/500 | 0.005 | 5.00E-07 | 1.00E-08 | | 35 | 1/500 | 0.05 | 5.00E-06 | 1.00E-08 |
| 6 | 1/2 | 0.005 | 5.00E-07 | 1.00E-07 | | 36 | 1/2 | 0.05 | 5.00E-06 | 1.00E-07 |
| 7 | 1/10 | 0.005 | 5.00E-07 | 1.00E-07 | | 37 | 1/10 | 0.05 | 5.00E-06 | 1.00E-07 |
| 8 | 1/50 | 0.005 | 5.00E-07 | 1.00E-07 | | 38 | 1/50 | 0.05 | 5.00E-06 | 1.00E-07 |
| 9 | 1/100 | 0.005 | 5.00E-07 | 1.00E-07 | | 39 | 1/100 | 0.05 | 5.00E-06 | 1.00E-07 |
| 10 | 1/500 | 0.005 | 5.00E-07 | 1.00E-07 | | 40 | 1/500 | 0.05 | 5.00E-06 | 1.00E-07 |
| 11 | 1/2 | 0.005 | 5.00E-07 | 1.00E-06 | | 41 | 1/2 | 0.05 | 5.00E-06 | 1.00E-06 |
| 12 | 1/10 | 0.005 | 5.00E-07 | 1.00E-06 | | 42 | 1/10 | 0.05 | 5.00E-06 | 1.00E-06 |
| 13 | 1/50 | 0.005 | 5.00E-07 | 1.00E-06 | | 43 | 1/50 | 0.05 | 5.00E-06 | 1.00E-06 |
| 14 | 1/100 | 0.005 | 5.00E-07 | 1.00E-06 | | 44 | 1/100 | 0.05 | 5.00E-06 | 1.00E-06 |
| 15 | 1/500 | 0.005 | 5.00E-07 | 1.00E-06 | | 45 | 1/500 | 0.05 | 5.00E-06 | 1.00E-06 |
| 16 | 1/2 | 0.01 | 1.00E-06 | 1.00E-08 | | 46 | 1/2 | 0.1 | 1.00E-05 | 1.00E-08 |
| 17 | 1/10 | 0.01 | 1.00E-06 | 1.00E-08 | | 47 | 1/10 | 0.1 | 1.00E-05 | 1.00E-08 |
| 18 | 1/50 | 0.01 | 1.00E-06 | 1.00E-08 | | 48 | 1/50 | 0.1 | 1.00E-05 | 1.00E-08 |
| 19 | 1/100 | 0.01 | 1.00E-06 | 1.00E-08 | | 49 | 1/100 | 0.1 | 1.00E-05 | 1.00E-08 |
| 20 | 1/500 | 0.01 | 1.00E-06 | 1.00E-08 | | 50 | 1/500 | 0.1 | 1.00E-05 | 1.00E-08 |
| 21 | 1/2 | 0.01 | 1.00E-06 | 1.00E-07 | | 51 | 1/2 | 0.1 | 1.00E-05 | 1.00E-07 |
| 22 | 1/10 | 0.01 | 1.00E-06 | 1.00E-07 | | 52 | 1/10 | 0.1 | 1.00E-05 | 1.00E-07 |
| 23 | 1/50 | 0.01 | 1.00E-06 | 1.00E-07 | | 53 | 1/50 | 0.1 | 1.00E-05 | 1.00E-07 |
| 24 | 1/100 | 0.01 | 1.00E-06 | 1.00E-07 | | 54 | 1/100 | 0.1 | 1.00E-05 | 1.00E-07 |
| 25 | 1/500 | 0.01 | 1.00E-06 | 1.00E-07 | | 55 | 1/500 | 0.1 | 1.00E-05 | 1.00E-07 |
| 26 | 1/2 | 0.01 | 1.00E-06 | 1.00E-06 | | 56 | 1/2 | 0.1 | 1.00E-05 | 1.00E-06 |
| 27 | 1/10 | 0.01 | 1.00E-06 | 1.00E-06 | | 57 | 1/10 | 0.1 | 1.00E-05 | 1.00E-06 |
| 28 | 1/50 | 0.01 | 1.00E-06 | 1.00E-06 | | 58 | 1/50 | 0.1 | 1.00E-05 | 1.00E-06 |
| 29 | 1/100 | 0.01 | 1.00E-06 | 1.00E-06 | | 59 | 1/100 | 0.1 | 1.00E-05 | 1.00E-06 |
| 30 | 1/500 | 0.01 | 1.00E-06 | 1.00E-06 | | 60 | 1/500 | 0.1 | 1.00E-05 | 1.00E-06 |

Table 1: Table containing all combinations of parameters for the supply of mutations, for most of the simulations in the results section (Figures 3 and 4).

| No. | description | parameter | standard value(s) |
|---|---|---|---|
| 1 | initial population size for species 1 and 2 | $N_1$, $N_2$ | 50, 50 |
| 2 | growth rates (initial value) | $r_1$,$r_2$ | varies in sim |
| 3 | intra-specific interaction coefficients (initial value) | $a_{11}$, $a_{22}$ | varies in sim |
| 4 | inter-specific interaction coefficients (constants) | $a_{12}$, $a_{21}$ | -0.000025, -0.000025 |
| 5 | proportion of trait-increasing mutations (both species) | $m_k$ | 1/500 |
| 6 | mutation rate (both species) | $\mu$ | $10^{-6}$ |
| 7 | multiplier for mutation effect size (both species) | $\beta_r$, $\beta_a$, | 0.05, 0.000005, |
| 8 | parameters for distribution of mutation effects (gamma distribution) | $s$, $\alpha$ | 1, 0.5 |
| 9 | constants in tradeoff function for species 1 ($a = f(r) = C_1 e^{C_2 * r}$) | $C_1$, $C_2$ | -0.0000206, 3.2 |
| 10 | constants in tradeoff function for species 2 ($a = f(r) = C_3 e^{C_4 * r}$) | $C_3$, $C_4$ | -0.0000108, 3.5 |
| 11 | end tick for simulation | $t_{\max}$ | 50,002 |
| 12 | log interval | $lt$ | 100 |

Table 2: Table containing parameters for two-species simulations with fixed values of mutation supply parameters but initial growth parameters varied (Figure 5).

| No. | description | parameter | standard value(s) |
|---|---|---|---|
| 1 | initial population size for species 1 and 2 | $N_1$, $N_2$ | 50, 50 |
| 2 | growth rates (initial value) | $r_1$,$r_2$ | 0.253579, 0.2035197 |
| 3 | intra-specific interaction coefficients (initial value) | $a_{11}$, $a_{22}$ | -0.000126, -0.0000511 |
| 4 | inter-specific interaction coefficients (constants) | $a_{12}$, $a_{21}$ | -0.000025, -0.000025 |
| 5 | proportion of trait-increasing mutations (both species) | $m_k$ | 1/500 |
| 6 | mutation rate (both species) | $\mu_1$, $\mu_2$ | varies between $10^{-7}$ and $10^{-6}$ |
| 7 | multiplier for mutation effect size (both species) | $\beta_r$, $\beta_a$, | 0.05, 0.000005, |
| 8 | parameters for distribution of mutation effects (gamma distribution) | $s$, $\alpha$ | 1, 0.5 |
| 9 | constants in tradeoff function for species 1 ($a = f(r) = C_1 e^{C_2 * r}$) | $C_1$, $C_2$ | -0.0000206, 3.2 |
| 10 | constants in tradeoff function for species 2 ($a = f(r) = C_3 e^{C_4 * r}$) | $C_3$, $C_4$ | -0.0000108, 3.5 |
| 11 | end tick for simulation | $t_{\max}$ | 50,002 |
| 12 | log interval | $lt$ | 100 |

Table 3: Table containing parameters for two-species simulations where mutation rates varied independently for both species (with all other parameters constant) (Figure 6).
